# Enhancing glycan occupancy of soluble HIV-1 envelope trimers to mimic the native viral spike

**DOI:** 10.1101/2020.07.02.184135

**Authors:** Ronald Derking, Joel D. Allen, Christopher A. Cottrell, Kwinten Sliepen, Gemma E. Seabright, Wen-Hsin Lee, Kimmo Rantalainen, Aleksandar Antanasijevic, Jeffrey Copps, Anila Yasmeen, Patricia van der Woude, Steven W. de Taeye, Tom L.G.M. van den Kerkhof, P.J. Klasse, Gabriel Ozorowski, Marit J. van Gils, John P. Moore, Andrew B. Ward, Max Crispin, Rogier W. Sanders

## Abstract

The HIV-1 envelope glycoprotein (Env) trimer is decorated with *N*-linked glycans, which are attached to asparagine residues in the amino acid sequon NxT/S by oligosaccharyltransferases (OST). Artificial glycan “holes” exist when a PNGS is under-occupied on recombinant Env-based vaccines, but not on their viral counterpart. Native-like SOSIP trimers, including clinical candidates, have these artificial holes in the glycan shield that induce strain-specific neutralizing antibodies (NAbs) or non-NAbs. To increase PNGS occupancy, eliminate artificial glycan holes, and mimic the glycosylation of native BG505 Env, we replaced all 12 NxS sequons on the BG505 SOSIP trimer with NxT, thereby increasing the affinity of the sequons for OST. All PNGS, except N133 and N160, were nearly fully occupied on the modified trimer. Occupancy of the N133 site could be increased by changing N133 to NxS, while occupancy of the N160 site could be restored by reverting the nearby N156 sequon to NxS. Hence, OST affinity can influence glycan occupancy when two PNGS are in close proximity. Increasing glycan occupancy should reduce off-target immune responses to artificial glycan holes on vaccine antigens.

## Introduction

The HIV-1 envelope glycoprotein spike (Env) initiates viral entry in host cells and is the only target for neutralizing antibodies (NAbs). Env is a trimer of heterodimers consisting of gp120 and gp41 subunits. During the early stages of protein synthesis Env is decorated by *N*-linked glycans which, upon leaving the Golgi apparatus, comprise approximately 50% of the total mass ^1–3^. The *N*-linked glycans play a critical role in various stages of the viral life cycle, including Env folding, binding to lectin receptors, and immune evasion by shielding underlying protein epitopes ^4,5^. The gp120 subunit can contain up to 35 potential *N*-linked glycosylation sites (PNGS) and gp41 contains typically 4 PNGS. Hence, there can be as many as 120 PNGS per trimer. Most of these PNGS are attachment points for *N*-linked glycans by the host cell glycosylation machinery, but some may remain unoccupied ^6^.

*N*-linked glycans are attached to asparagine residues (N) located in NxT/S sequons where T is threonine, S is serine and x can be any amino acid except proline. The amino acid that follows the NxT/S sequon can also affect glycan attachment: a proline at this position abrogates glycosylation^7^. The enzyme responsible for glycan attachment is oligosaccharyltransferase (OST), which consists of seven to eight non-identical subunits. The catalytic subunit has two isoforms STT3A and STT3B ^8,9^. The STT3A subunit is associated with the translocon and attaches glycans during translation, but it sometimes skips sequons that are in close proximity, i.e. NxT/S-NxT/S or NxT/S-x-NxT/S motifs (termed gap-0 and gap-1 sites, respectively). The STT3B subunit facilitates co-translational and post-translational glycosylation of sequons that are skipped by STT3A. Even so, not all gap-0 and gap-1 sites are fully occupied by glycans ^9^. HIV-1 Env contains a number of gap-0 and gap-1 sites, i.e. the N156/N160 gap-1 site that contains glycans that are essential for several broadly neutralizing antibodies (bNAb) (Los Alamos database; http://www.hiv.lanl.gov). ^9–12^.

The protein surface of Env is almost completely shielded by glycans, which contributes to its poor immunogenicity ^13^. Env and Env-based vaccines, including native-like SOSIP trimers and SOSIP-based clinical candidates, have holes in their glycan shield from two different sources.

Firstly, a conserved PNGS can be absent from the Env of any particular isolate; for example, the conserved glycosylation sequons at N241 and N289 are not present in the BG505 virus and SOSIP trimers. The absence of these two glycans creates a large immunodominant hole in the glycan shield that is responsible for generating the majority of strain-specific NAb responses induced by BG505 SOSIP trimers in animals ^14,15^. Of note is that HIV-1 isolates with an intact glycan shield are more capable of inducing broader neutralizing responses than ones with multiple holes, which may act as immunological decoys and thereby impede the development of neutralization breadth ^16^. Filling glycan holes may therefore be a strategy to focus the immune response on more desirable targets; it can be achieved simply by restoring the missing PNGS ^17,18^.

Secondly, a PNGS that is in the protein sequence can be under-occupied, probably because the site is often skipped by STT3A and STT3B ^8,9,19^. Several PNGS in the V1/V2 domain of BG505 SOSIP.664 trimers are under-occupied, creating artificial glycan holes ^20^. The PNGS at position 611 of these trimers is probably also under-occupied, because it induces antibodies that can only neutralize viruses from which N611 is deleted ^14,15,20,21^. Direct comparisons revealed that glycan occupancy is lower on BG505 SOSIP trimers than on the sequence-matched viral Env ^20,22^. However, increasing the occupancy of a PNGS that is caused by OST skipping is not straightforward because the factors that determine whether a PNGS becomes a substrate for OST are poorly understood. Studies on model glycoproteins have, however, shown that NxT sequons have a two- to three-fold higher probability of becoming glycosylated compared to NxS ^23^, most likely because OST has a higher affinity for NxT ^24,25^. In this study, we sought to increase PNGS occupancy on prototypic BG505 SOSIP trimers to an extent that mimicked viral BG505 Env, and thereby eliminate artificial glycan holes.

## Results

### Cryo-EM confirms N611 under-occupancy

The glycan compositions of several native-like HIV-1 Env trimers have been characterized previously ^26–32^. PNGS occupancy analyses of soluble trimers and their viral counterparts are less common, but new techniques have enabled such studies on BG505 SOSIP.664 and other trimers ^20,22,33,34^. Here, we performed cryo-electron microscopy (cryo-EM), PNGS occupancy and glycan composition analyses on BG505 SOSIP.v4.1 trimers (from here on referred to as WT proteins) ^35^. Monoclonal antibodies (MAbs) have been isolated from rabbits and rhesus macaques that were immunized with BG505 SOSIP.664 trimers by single B-cell sorting and B-cell receptor (BCR) cloning ^14,15,21^. While many of the MAbs targeted the N241/N289 glycan hole, a subset of MAbs were directed against the N611 site. These MAbs could bind the WT proteins, but were unable to neutralize the corresponding virus unless the N611 PNGS was knocked-out ^14,15,20,21^. The implications of these findings are that the region underneath the N611 glycan on BG505 SOSIP trimers is immunogenic for MAb induction because the N611 PNGS is under-occupied; and that the same site is more extensively occupied on the MAb-resistant virus.

To probe the N611 underoccupancy on the BG505 SOSIP trimer, we used MAb RM20E1, directed to the N611 site that was isolated from a BG505 SOSIP.664-immunized macaque and performed cryo-EM studies with the Fab in complex with the BG505 SOSIP.v5.2 trimer. ^14,21,36^. Multiple rounds of 3D sorting were performed to segregate trimers with zero, one, two, or three RM20E1 Fabs attached (Fig 1A). Density corresponding to the N611 glycan was observed on all protomers with no Fab attached, but in contrast there was no visible N611 density on any protomers bound to RM20E1 (Fig. 1B). These observations are consistent with the previously described binding and neutralization results ^15,21^. A further analysis of the numbers of trimers in 3D-sorted classes with zero, one, two, or three RM20E1 Fabs showed that N611 was occupied on ∼40% of the protomers.

**Fig. 1.**
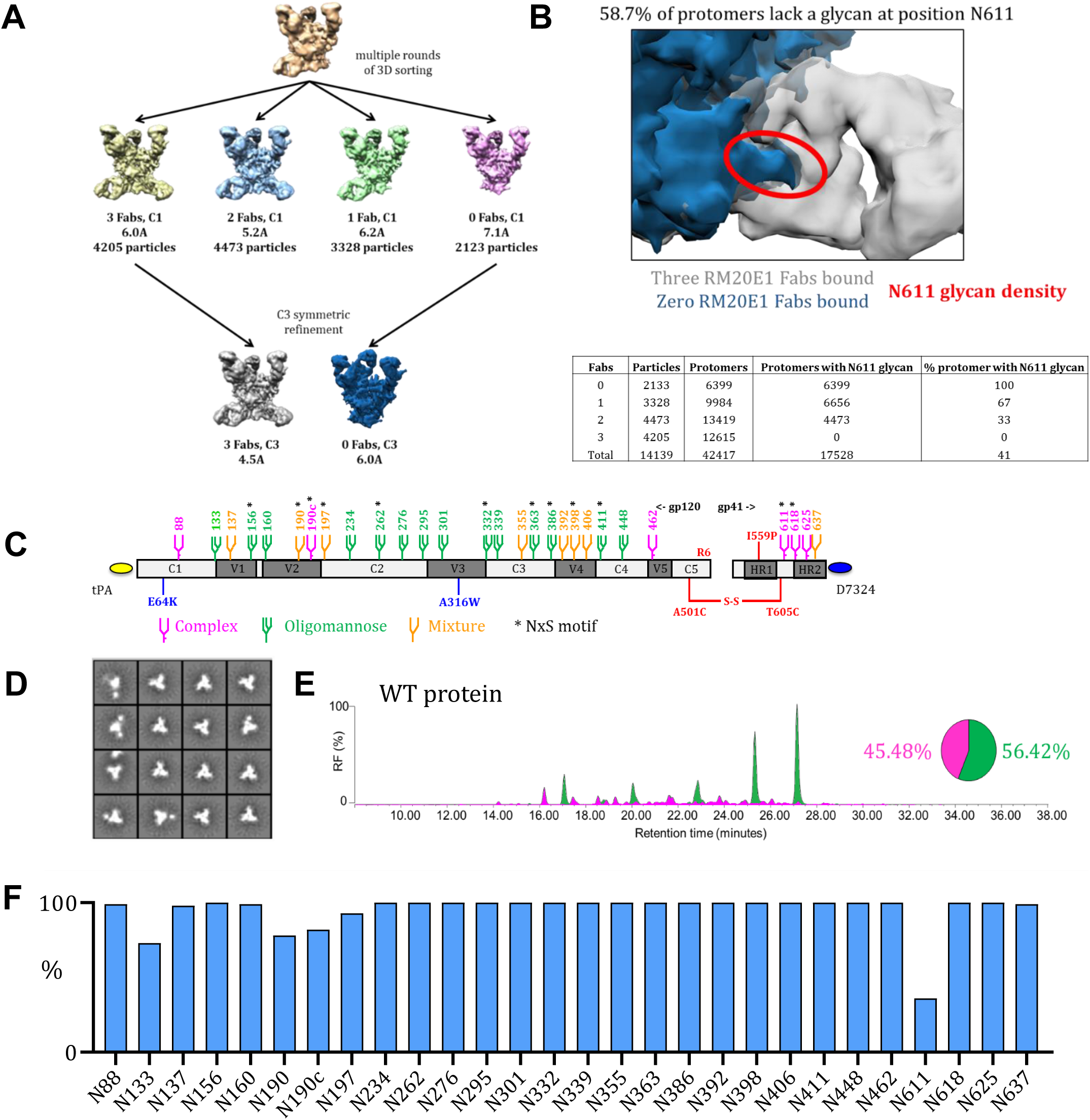
Several PNGS on BG505 SOSIP.v4.1 trimers are under-occupied. **(A)** Cryo-EM analysis of WT proteins in complex with the RM20E1 Fab. Complexes were sorted based on the number of Fabs bound; the numbers of particles in each reconstruction are listed, as are the resolutions of the final reconstructions. **(B)** Detail of an overlay of the Cryo-EM reconstructions of the BG505 SOSIP.664 trimer alone (in blue) and in complex with the RM20E1 Fab (in grey). The density of the N611 glycan (also in blue) on the trimer without RM20E1 is highlighted by the red oval to illustrate its clash with the RM20E1 Fab (in grey). The total number of complexes with different numbers of Fab bound are indicated in the table, as are the protomers with N611 glycan density. This density was only observed on protomers that did not bind the RM20E1 Fab. From these numbers, the % of protomers that lack the N611 glycan could be calculated (∼59%). **(C)** Linear representation of the D7324-tagged BG505 SOSIP.v4.1 trimer. The SOSIP changes and the stabilizing E64K and A316W mutations are all highlighted ^35,38^. The glycans are also indicated, with the amino acid numbering based on the HXB2 strain. Sites that are predominantly occupied by oligomannose species are colored green and by complex species in magenta, while orange indicates that either an oligomannose or a complex glycoform can be present, based on data presented in Fig. 1D. An asterisk indicates that the PNGS is encoded by an NxS sequon. The individual sequon compositions are in Table S1. **(D)** NS-EM analysis of WT proteins, showing the 2D class-averages. **(E)** HILIC-UPLC analysis of WT proteins. Peaks colored green represent glycans that can be cleaved by endoglycosidase H (endoH) and correspond to oligomannose or hybrid-type glycans. The HILIC-UPLC spectra and pie chart show the overall oligomannose glycan (green) and complex/hybrid glycan (magenta) content. **(F)** Quantification of site-specific occupancy for all 28 PNGS on WT proteins, derived from LC-ESI MS analyses. Results are the mean of two independent biological replicates of the WT protein. All the data in panels A-F were derived using WT proteins produced in HEK293F cells followed by PGT145-affinity purification. The corresponding data on WT proteins purified by the 2G12/SEC method are in Fig. S1.

### Several PNGS on BG505 SOSIP.v4.1 trimers are under-occupied

Next, we used several techniques to gain a more detailed understanding of the glycan shield on the WT protein. The D7324-tagged WT protein (Fig. 1C) was expressed in HEK293F cells and purified by affinity chromatography using the PGT145 bNAb, which is selective for native-like trimers ^37^. We also affinity purified the D7324-tagged WT protein using the 2G12 bNAb followed by size exclusion chromatography (SEC) ^35,38^. Both methods are routinely used for purifying native-like trimers, including in cGMP programs ^37,39^. WT proteins purified via the PGT145 and 2G12/SEC methods had a homogeneous native-like conformation as judged by negative stain electron microscopy (NS-EM) (Fig. 1D; Fig. S1A). Overall, analyses of the PGT145- and 2G12/SEC purified-proteins yielded very similar data, with a few notable exceptions (see below). We present data on the PGT145-purified proteins in the main Figures, with the corresponding Figures for their 2G12/SEC-purified counterparts in Supplemental Information.

To determine the total oligomannose content of the purified WT protein, we performed hydrophilic interaction liquid chromatography-ultra performance liquid chromatography (HILIC-UPLC) ^20^. The PGT145-purified WT protein contained 56% oligomannose-type glycans and 60% when purified using 2G12/SEC. In both cases, Man_8_GlcNAc_2_ and Man_9_GlcNAc_2_ were the predominant glycan species (Fig. 1E; Fig. S1B), consistent with previous reports on native-like SOSIP trimers ^20,31,35^.

To investigate PNGS occupancy and glycan composition, we performed a site-specific analysis of the WT protein using liquid chromatography-electrospray ionization (LC-ESI) mass spectrometry (MS) on an Orbitrap Fusion mass spectrometer (Thermo Fisher Scientific) ^20,31^ (Fig. 1F). PNGS occupancy is expressed as the percentage of the total peptide that is modified by a glycan; we considered a modification level of ≥ 95% as indicative of full occupancy. The majority of PNGS on gp120 from the PGT145-purified WT protein were fully occupied, including the five located within a 25-amino acid stretch of V4 (N386, N392, N398, N406, and N411). However, under-occupancy was seen for sites near the apex: N133: ∼75% occupancy; N190: ∼80%; N190c: ∼80%; and N197: ∼93% (Fig. 1F). The same sites were also under-occupied on the 2G12/SEC-purified WT protein: N133: ∼90% occupancy, N137: ∼80%, N190: ∼90%; and N190c: ∼80% (Fig. S1C). These regions of under-occupancy are similar to those described previously for BG505 SOSIP.664 trimers ^20^.

In our previous study, we did not perform a site-specific PNGS occupancy analysis for gp41 ^20^. We now report that the N611 glycan on the PGT145-purified protein is markedly under-occupied (∼35% occupancy) (Fig. 1F; Fig. S1C). This finding is highly concordant with the cryo-EM analysis (Fig. 1A). The other three gp41 sites (N618, N625 and N637) were fully occupied on the PGT145-purified WT protein. However, the N625 site was less occupied (∼70%) and the N611 site more occupied (70%) on the 2G12/SEC-purified WT protein (Fig. S1C).

The LC-ESI MS method also allowed us to determine the specific glycoforms present at each site. For most PNGS on the WT protein, the glycoforms were very similar to those observed in our previous studies ^20,31^ (Fig. S1D). Thus, sites N156, N160, N234, N262, N276, N295, N332, N339, N363, N386, N392 N411 and N448 were predominantly occupied by oligomannose glycans, sites N190c, N398, N611, N618 and N625 by complex glycans, and sites N88, N137, N190, N197, N301, N355, N406, N462 and N637 by a mixture of both forms.

### Glycan occupancy is enhanced by PNGS sequon engineering

Although the determinants of glycan occupancy are not precisely clear, studies on model glycoproteins have shown that the NxT sequon has a two-to three-fold higher probability of becoming glycosylated compared to an NxS sequon ^23^. The relative use of NxS (∼40%) *versus* NxT (∼60%) in the BG505 sequence is consistent with most isolates in the Los Alamos database (Table S1). Thus, when an NxS sequon is usually present at a particular site, that is also the case for BG505. To explore how to increase glycan occupancy, we first designed two stabilized BG505 SOSIP.v4.1 trimers ^35^ in which we changed all the PNGS to either NxT or NxS. The two variants are designated NxT protein and NxS protein, respectively.

We used SDS-PAGE and Blue Native (BN-) PAGE to assess the expression and trimer formation of unpurified WT, NxT and NxS proteins produced in transfected HEK293T cells (Fig. S2A and B). The NxT protein was expressed efficiently and formed trimers, although slightly less well than the WT protein, but the NxS protein was not expressed efficiently. A D7324-capture ELISA using bNAbs 2G12, VRC01, PGT145 and PGT151 to probe the folding of the proteins in the unpurified supernatants confirmed these findings (Fig. S2C). Thus, all four bNAbs bound to the NxT and WT proteins, but the NxS protein could only be detected using 2G12. The defect in NxS protein expression might be caused by the absence of glycans that are important for Env to fold properly in the ER ^40^.

Purified, HEK293F cell-expressed NxT proteins were fully cleaved with no evidence of aggregation or dissociation when analyzed by non-reducing and reducing SDS-PAGE or BN-PAGE (Fig. S2D). Compared to WT, the NxT protein migrated slightly more slowly through the SDS- and BN-PAGE gels, which is suggestive of more extensive glycosylation. The NS-EM 2D-class averages showed that the NxT protein was in a native-like conformation (Fig. 2A, Fig. S3A).

**Fig. 2.**
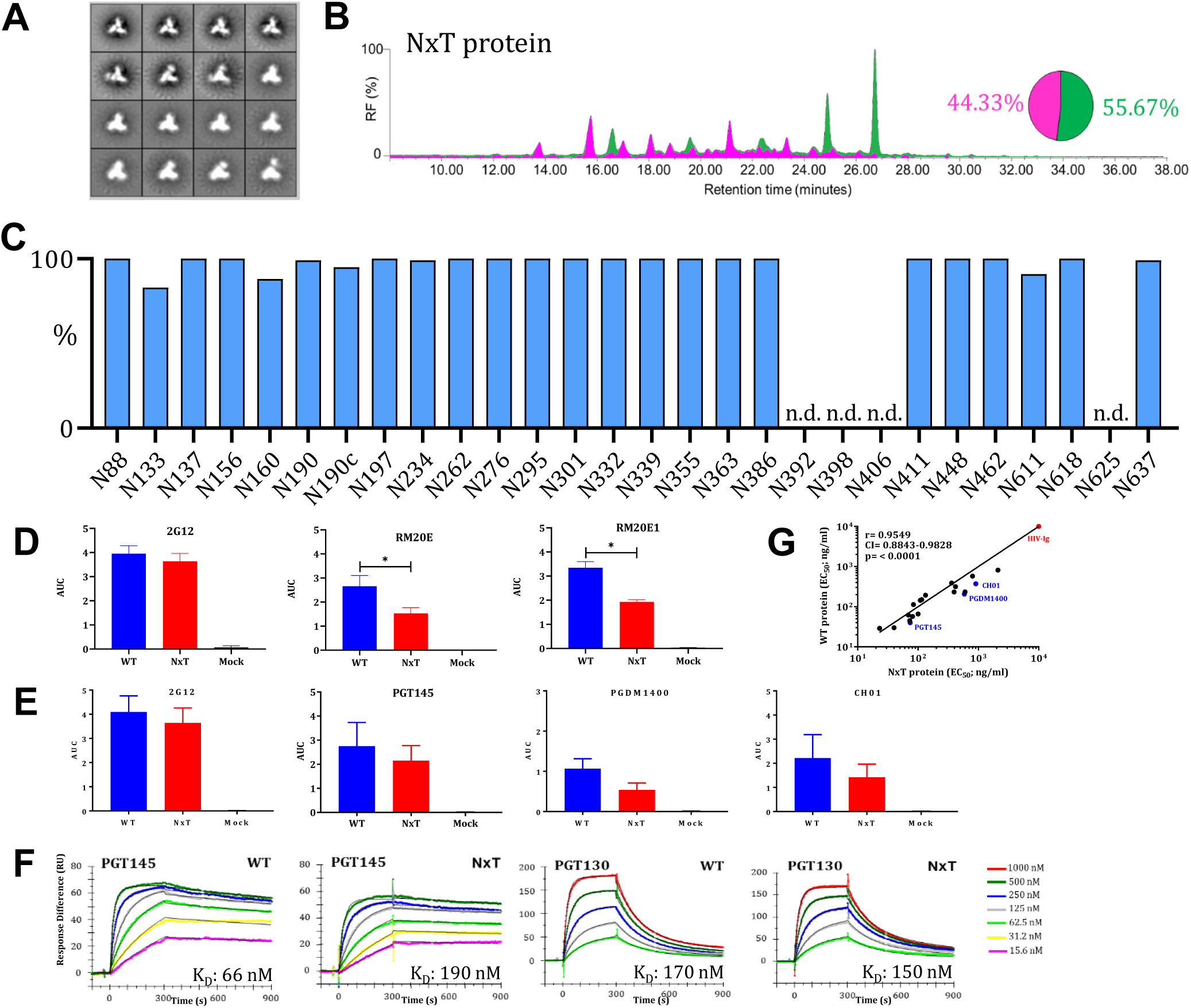
Glycan occupancy is increased by PNGS sequon engineering. **(A)** NS-EM analysis of NxT proteins, showing the 2D class-averages. **(B)** HILIC-UPLC analysis of the NxT protein. The color coding of the spectra and pie chart is the same as in Fig. 1E. **(C)** Quantification of site-specific occupancy for all 28 PNGS on NxT proteins, derived from LC-ESI MS analyses. **(D)** Binding of WT and NxT proteins to two N611-directed non-NAbs, RM20E and RM20E1, isolated from BG505 SOSIP.664 immunized rhesus macaques. The area under the curve (AUC) values derived from ELISA titration curves are plotted. * indicates a significant difference (P <0.05) between the WT and NxT proteins, calculated using a Mann-Whitney 2-tailed test. **(E)** Binding of WT and NxT proteins to three V2-apex directed bNAbs, PGT145, PGDM1400 and CH01, and also 2G12 for comparison. AUC values derived from derived from ELISA titration curves are plotted. **(F)** Binding of bNAbs PGT145 and PGT130 to WT and NxT proteins, assessed by SPR. A summary of the binding kinetics for both bNAbs is in Fig. S3E. **(G)** The EC_50_ values derived using WT and NxT proteins were plotted and compared using Spearman’s correlation coefficient, r, and Prism software version 7.03. All analyses were performed on NxT proteins produced in HEK293F followed by PGT145 purification. The corresponding data on NxT proteins purified by the 2G12/SEC method are in Fig. S3.

The overall glycan composition of the NxT protein was very similar to WT, with oligomannose contents of 56% and 60% after PGT145 and 2G12/SEC-purification, respectively (Fig. 2B; Fig. S3B). However, the LC-ESI MS experiments revealed some striking differences in PNGS occupancy. Whereas the N190, N190c and N197 PNGS were under-occupied on the WT protein (Fig. 1F), the same sites were >95% occupied on both preparations of the NxT proteins (Fig. 2C; Fig S3C). Furthermore, occupancy of N611 on gp41 increased from ∼40% and ∼70% on the PGT145- and 2G12/SEC-purified WT proteins, respectively, to ∼90% and ∼85% for the corresponding NxT proteins (Fig. 2C; Fig S3C).

Occupancy at N133 was not adequately restored on the PGT145-purified NxT protein as it increased from 73% to 83%, which means that ∼1 in 5 protomers still lack a glycan at this position. Conversely, N133 occupancy actually decreased to ∼60% on the 2G12/SEC-purified NxT protein, compared to ∼95% for WT (Fig. S3C). We are unable to explain this unexpected finding. Furthermore, while the N160 PNGS was fully occupied on the PGT145-purified WT protein, that was not the case on the NxT protein as occupancy decreased to ∼85% (Fig. 2C). An even greater decrease in N160 occupancy, to ∼70%, was seen for the 2G12/SEC-purified NxT protein (Fig. S3C). The PGT145 epitope requires at least two occupied N160 sites, which probably explains the higher N160 occupancy on the PGT145-*versus* 2G12/SEC-purified NxT protein^12^. The decreased N160 occupancy on the NxT protein *versus* WT is studied further below.

There were no substantial differences when the glycoforms at each PNGS of the NxT protein were compared to the WT protein. The sites that were more occupied on the NxT protein than on WT (N190, N190c, N197 and N611) were all predominantly occupied by complex glycans (Fig. S3C).

The increased occupancy of N611 on NxT proteins could reduce the immunogenicity of non-NAb epitopes associated with this artificial glycan hole. To assess this *in vitro*, we used two N611-directed non-NAbs isolated from BG505 SOSIP.664-vaccinated macaques ^38^. Using a D7324-capture ELISA, the PGT145-purified NxT protein was significantly less reactive than the WT protein with non-NAbs RM20E (P=0.03) and RM20E1 (P=0.03), calculated using a two-tailed Mann-Whitney test (Fig. 2D).

The overall antigenicity of the PGT145-purified NxT proteins was assessed by ELISA (Fig. S4). Most of the bNAbs tested recognize epitopes involving glycans; they include ones directed against quaternary-structure dependent V1/V2-apex sites (PG9, PG16, PGT145, PGDM1400 and CH01), the N332-glycan dependent epitope cluster (PGT121-123, PGT125-128, PGT130, 2G12 and PGT135), the quaternary-structure dependent epitopes at the gp120-g41 interface (PGT151 and 35O22), and gp41 (3BC315). We also used bNAbs against the CD4bs (VRC01 and 3BNC60) and polyclonal HIV-Ig. In all cases, the bNAbs bound similarly to the WT and NxT proteins, with <3-fold difference in 50% binding concentrations (EC_50_) (Fig S4). An area under the curve (AUC) analysis, however, revealed that bNAbs against the trimer apex were less reactive with the NxT protein (Fig. 2E). This finding is concordant the reduced occupancy of N160 on this trimer, as the epitopes involve this glycan. Similar results were obtained when the 2G12/SEC-purified WT and NxT proteins was tested by ELISA (Fig. S3D), and by surface plasmon resonance (SPR) (WT, K_D_ 66 nM *versus* NxT, K_D_ 190 nM for PGT145; the K_D_ values are for the initial interaction as fitted with a conformational-change model _41_) (Fig. 2F; Fig.S3E). The EC_50_ values for bNAb binding to the WT and NxT proteins were highly correlated (Spearman r=0.95, P<0.0001 (Fig. 2G) and Spearman r=0.97, P<0.0001 (Fig. S3F)).

To verify whether NxT sequon engineering was compatible with Env function, we generated a BG505.T332N infectious molecular clone (IMC) in which we changed all Env PNGS to NxT. The NxT and WT viruses were equally infectious (Fig. S5A). In contrast, an IMC with the PNGS all changed to NxS was not infectious, which is consistent with the poor expression of the corresponding NxS protein (Fig. S2; Fig. S5A). In a neutralization assay, the NxT virus had reduced sensitivity to PGT145 (26-fold increase in IC_50_; Fig. S5C and D). A second V1/V2-apex bNAb, CH01, neutralized the NxT and WT viruses comparably, as judged by IC_50_ values, but the maximum extent of neutralization was only ∼60% for the NxT virus while its WT counterpart was completely neutralized at high CH01 concentrations. (Fig. S5C). The impact of the NxT modification on the V1/V2-apex are therefore seen with viruses and recombinant trimers, and are consistent with under-occupancy of the PNGS at N160 in both contexts.

### Occupancy of gap-1 sites can be enhanced by reducing OST affinity of the first site

The reduced N160 occupancy on the NxT protein was surprising since the glycan sequon was unchanged compared to WT. However, the preceding N156 sequon was changed from NxS to NxT. This site is fully occupied on WT proteins purified by both methods (Fig. 1F; Fig. S1D). We hypothesized that increasing the affinity of the PNGS at N156 for OST by changing it to NxT might interfere with glycan attachment at N160. We therefore reverted T158 to the original S in the background of the NxT protein. The purified NxT T158S protein adopted a native-like conformation (Fig. 3A; Fig. S6A) and had an overall high oligomannose content of 50% and 48% for PGT145- and 2G12/SEC-purified proteins, respectively (Fig. 3B; Fig S6B). Of note is that the T158S mutation enhanced N160 occupancy to >95% for both preparations of purified proteins (Fig. 3C; Fig. S6C). That high occupancy level is now similar to what is seen for N160 on the WT protein (Fig. 1D; Fig. S1D).

**Fig. 3.**
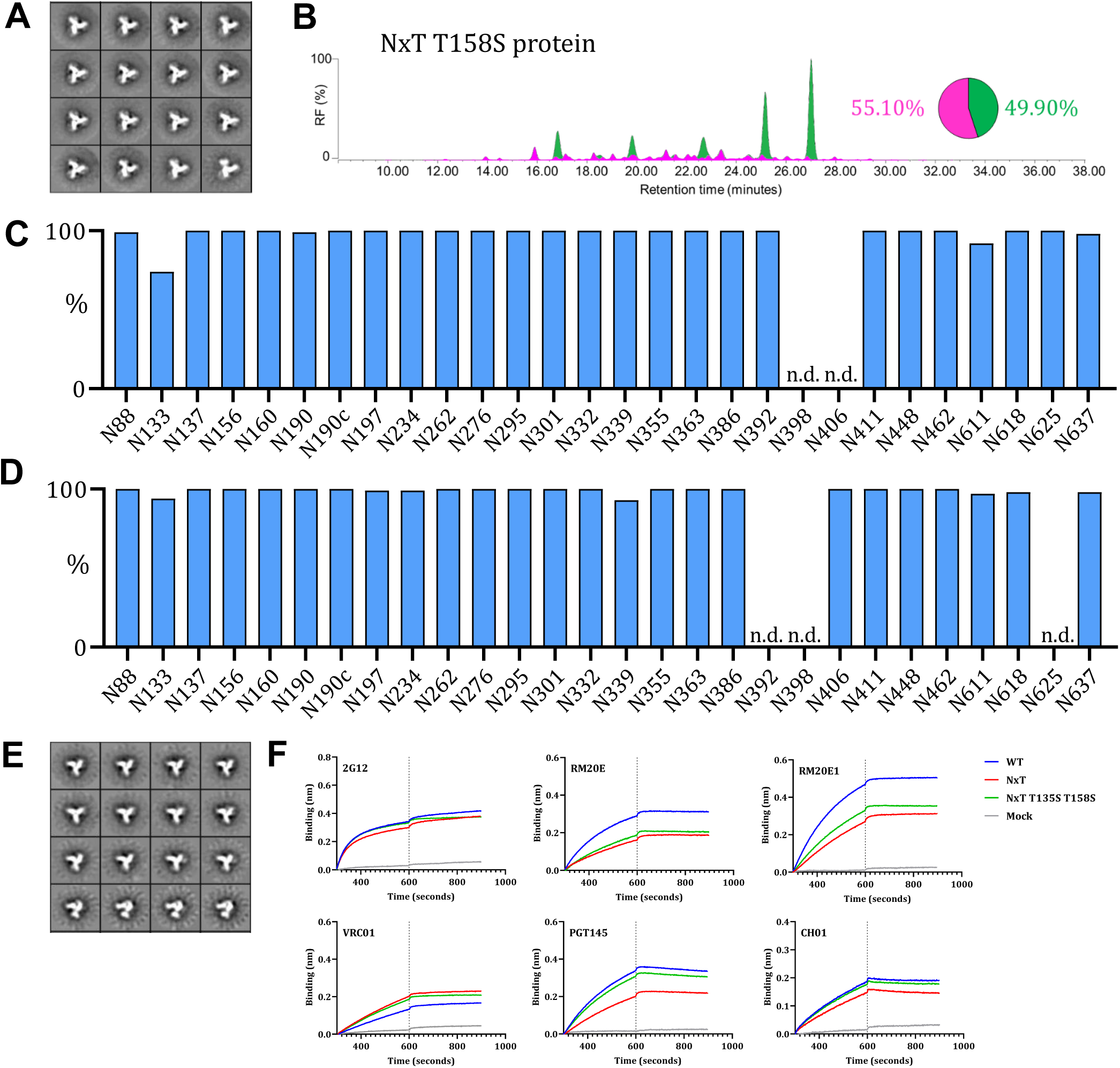
Occupancy of gap-1 sites can be increased by reducing the affinity of the first site for OST. **(A)** NS-EM analysis of NxT T158S proteins, showing the 2D class-averages. **(B)** HILIC-UPLC analysis of the NxT T158S protein. The color coding of the spectra and pie chart is the same as in Fig. 1E. **(C)** Quantification of glycan occupancy using LC-ESI MS of the 28 PNGS on NxT T158S proteins. **(D)** Quantification of site-specific occupancy for all 28 PNGS on NxT T135S T158S proteins, derived from LC-ESI MS analyses. Results shown are the mean of two independent biological replicates of the NxT T135S T158S protein. The corresponding data on NxT T135S T158S proteins purified by the 2G12/SEC method are in Fig. S6. **(E)** NS-EM analysis of NxT T135S T158S proteins, showing the 2D class-averages. **(F)** Binding of non-NAbs RM20E and RM20E1 to WT, NxT and NxT T135S T158S proteins, assessed by BLI. The bNAb 2G12 was also tested, for comparison. We also tested the binding of bNAbs VRC01, PGT145 and CH01. The average binding curves from 3 independent duplicates are shown. All analyses were performed on NxT T135S T158S trimers produced in HEK293F cells and affinity purified using PGT145.

The only site that remained considerably under-occupied on the NxT T158S protein was N133 (∼75% occupancy; Fig. 3C). Based on our success with the N160 site, and noting that the N133 and N137 sequons are spaced similarly to N156 and N160, we adopted the same strategy to increase N133 occupancy. Accordingly, we changed residue 135 from T to S in the background of the NxT T158S protein. The outcome was that on the PGT145-purified protein, occupancy of the N133 site increased from ∼75% for the NxT T158S protein (Fig. 3C) to ∼95% on NxT T135S T158S (Fig. 3D, E; Fig. S7A). Every PNGS was now nearly fully (>95%) occupied on the NxT T135S T158S protein (almost all sites >95% occupied) (Fig. 3D). However, for 2G12/SEC-purified NxT T135S T158S, N133 and N190 occupancy remained lower (∼65% and 86%, respectively (Fig. S7B)). Hence, the affinity purification method can impact PNGS occupancy.

The glycoforms at each site on the PGT145-purified NxT T135S T158S and WT proteins were very similar, but with a few exceptions. The N88, N190 and N406 glycans were more processed and closer resemble what is found on viral Env (Fig. S7B) ^20^. Conversely, the N133 glycan was less processed on this protein than WT. The glycans at N137, N276 and N406 were also more processed on the 2G12/SEC-purified NxT T135S T158S protein (Fig. S7B). The increased processing at N276 is similar to what was seen with the similarly purified NxT T158S protein, but differs from their PGT145-purified counterparts.

In a Biolayer interferometry (BLI) assay, MAbs RM20E and RM20E1 were less reactive with the PGT145-purified NxT T135S T158S protein than WT, which is further evidence that N611 is more occupied on the modified trimer (Fig. 3F). BLI also showed that the apex-targeting bNAbs PGT145 and CH01 were equally reactive with the WT and NxT T135S T158S proteins (Fig. 3F). Thus, increasing N160 occupancy via the T158S mutation improved the presentation of bNAb epitopes at the trimer apex.

In summary, the PGT145-purified NxT T135S T158S trimer closely mimics viral BG505 Env in respect of overall PNGS occupancy (Fig. 4). In particular, the artificial glycan hole recognized by non-NAbs and caused by under-occupancy of the N611 site is now closed ^20,22^, which could facilitate immune-focusing to more relevant neutralizing epitopes.

**Fig. 4.**
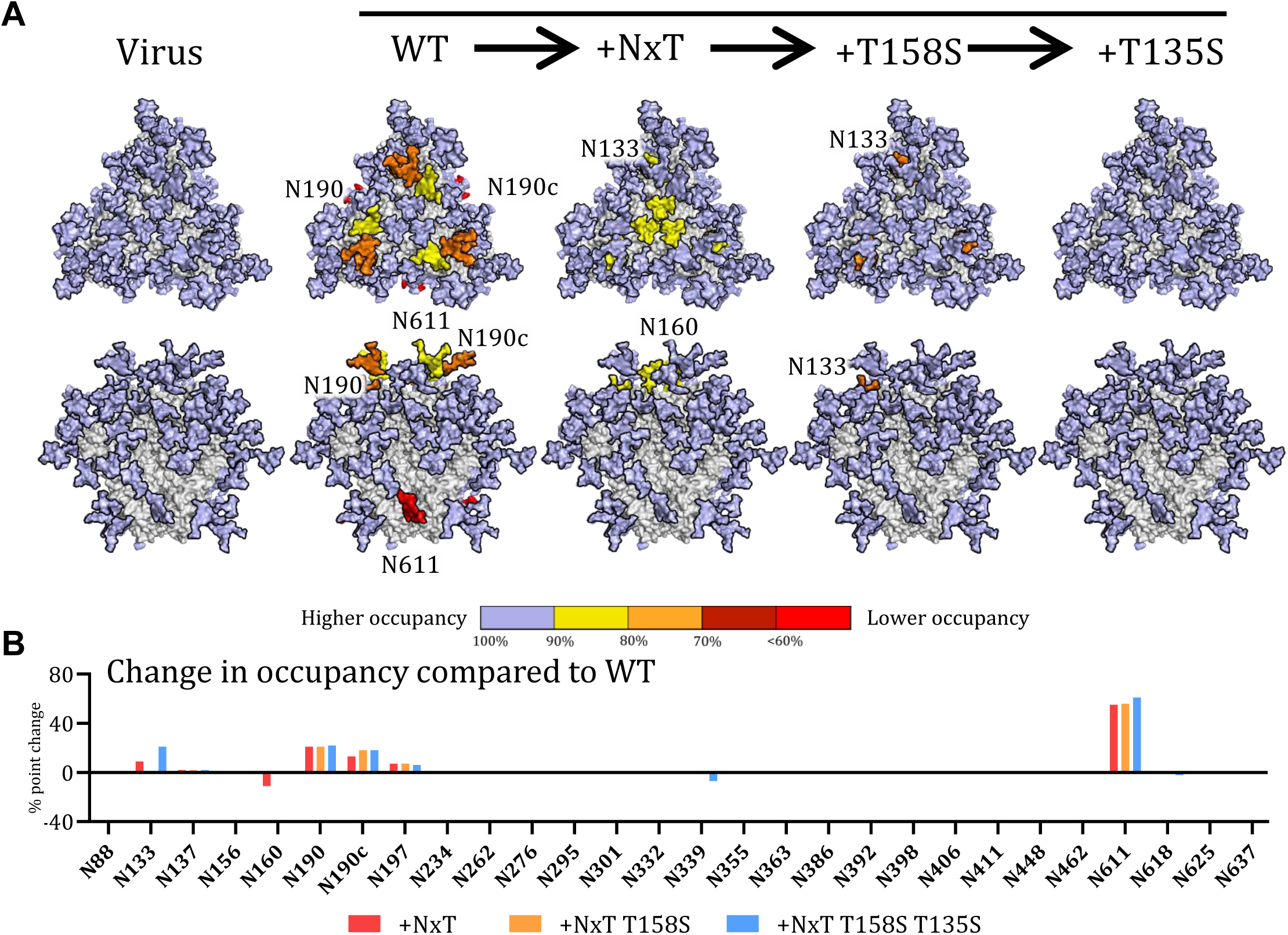
Sequential improvement of PNGS occupancy on SOSIP trimers. **(A)** Occupancy of each PNGS on the sequentially improved SOSIP.v4.1 trimers (WT, NxT, NxT T158S, NxT T135S T158S) is compared with the same site on the BG505 virus. Glycans are modelled onto each PNGS on a 3D model of the WT protein. Color coding reflects the occupancy of each PNGS: light blue, full occupancy (>95%); yellow, 80-95% occupancy; orange, 70-80% occupancy, dark red, 60-70% occupancy; bright red, <60% occupancy. **(B**) The sequential changes in PNGS occupancy at each PNGS are shown by the bars. The data shown are the percentage changes in occupancy caused by the various PNGS sequon changes, compared to the same site on the WT BG505 SOSIP.664 trimer.

## Discussion

In this study, we sought to overcome the under-occupancy of some PNGS on soluble BG505 SOSIP trimers and establish guidelines for similar efforts on other vaccine antigens. The absence of a PNGS can easily be resolved by re-introducing the site, which has been successfully achieved for BG505 SOSIP trimers ^17^. Under-occupancy of a PNGS is less easily addressed, but we were able to increase occupancy on these trimers by manipulating the affinity of PNGS sequons for OST. In our final construct, 11 of the 12 NxS sequons were changed to NxT while, conversely, 1 of the 16 NxT sequons was switched to NxS. Nearly all of the PNGS were fully occupied on the resulting BG505 SOSIP NxT T135S T158S trimer, which now better resembled the viral BG505 trimer in this regard. The modified SOSIP trimer should be a suitable design platform for further immune-focusing efforts intended to facilitate the induction of bNAbs, although additional efforts should be made to close the large glycan hole present at the trimer base ^42^.

PNGS under-occupancy occurs when the catalytic subunits of OST (STT3A and STT3B) skip sites, which occurs relatively frequently when the PNGS are close together (gap-0 or gap-1 sites) ^9^. The WT protein was under-occupied at N133 (gap-1), N190 (gap-0), N190c (gap-0), N197, and N611. In contrast, these same sites are fully occupied on viral Env ^20,22^. Several factors associated with recombinant protein expression may be relevant to the differences. First, viral Env is tethered to the ER membrane via its transmembrane domain (in addition to the signal peptide) which might promote association with OST, particularly for PNGS near the C-terminus of the nascent polypeptide. The dwell time in the ER may also be affected resulting in more or less time for OST to act ^22^. Second, the tissue plasminogen activator (TPA) signal peptide used to express the WT protein, which is cleaved off co-translationally, might impact interactions with OST differently than for the WT HIV-1 signal peptide, which is cleaved off post-translationally ^43,44^. Third, codon optimization of the recombinant SOSIP trimer may play a role via increased expression levels ^45^. The much lower expression of viral Env compared to its recombinant counterpart could cause sequon skipping by OST because of low substrate abundance. Fourth, the use of different producer cell lines (HEK239F or CHO cells for SOSIP trimers, CD4+ T cells for BG505 virus ^20,22,33,46^) may be relevant because of variation in their levels of OST and other components of the glycoprotein synthesis machinery.

Whatever the reasons for the under-occupancy of sites on the WT protein, we were able to address them. NxT sequons are more efficiently glycosylated than NxS, probably because their affinity for OST is higher. The preference of OST for T over S at the +2 position of PNGS may arise from stronger van der Waals interactions ^24,25^. Changing all 12 NxS sequons to NxT was compatible with efficient Env expression and trimer formation. When applied to the BG505 virus, infectivity was unaffected, a sensitive indicator that the 12 amino acid substitutions had no major adverse effects. PNGS occupancy on the NxT protein was increased at three sites surrounding the trimer apex, N190, N190c and N197, to the extent that they were now fully occupied (Fig. 4). The greatest change involved the N611 glycan on gp41, where occupancy more than doubled (i.e., from ∼40% to ∼90% occupancy). The outcome was markedly lower binding of N611-directed non-NAbs induced by immunization with BG505 SOSIP trimers that contained an artificial hole in the glycan shield at this position. Changing the N611 sequon from NxS to NxT thus effectively filled an immunodominant non-NAb glycan hole. Finally, PGT145-purification also selected for higher occupancy variants at the far distant N625 site, for reasons we do not yet understand.

In contrast to the increases seen for N190, N190c, N197 and N611, occupancy of N160 on the NxT protein was unexpectedly decreased, even though the N160 sequon itself was unchanged. The reduced occupancy of N160 diminished the binding of several V1/V2 bNAbs to the NxT protein, because their epitopes involve this glycan. Neutralization of the NxT virus by the same bNAbs was also substantially reduced compared to the WT virus. Here, the strongest effect was seen with PGT145, which was 26-fold less potent based on IC_50_ comparisons; a subtler difference was observed with CH01, as its maximum extent of neutralization was reduced from 100% for the WT virus to 60% for the NxT variant.

The analysis of N160 occupancy revealed the value of using the PGT145-purification method over 2G12/SEC. Thus, N160 under-occupancy was more pronounced on 2G12/SEC-purified NxT trimers, probably because the PGT145-affinity column selectively enriches a population of trimers that bind this bNAb strongly. PGT145 engages two N160 glycans and hence will positively select trimers with two or three glycans while largely ignoring the sub-populations with one or no glycan at this position ^12^. Accordingly, only a small proportion of PGT145-purified trimers contains unoccupied N160 sites. We found that PGT145 selected for fully occupied N133 and N137 PNGS for reasons we do not fully understand, although we note that these glycans are close to the PGT145 epitope ^12^.

Occupancy of N160 on the NxT protein could be restored fully by changing the N156 sequon back to NxS. Thus, decreasing the affinity of the N156 sequon for OST increased the efficiency of glycan addition at the downstream N160 site. As the N156 site is already fully occupied on WT proteins, the change to NxS could not further increase its occupancy. All the PNGS on the resulting NxT T158S protein were now fully occupied, except for N133 (∼75% occupancy). The N133 sequon is already in the apparently optimal NxT form in BG505 and most other HIV-1 Env sequences. However, and paradoxically, N133 occupancy could be improved by reducing OST affinity at this site via a change to NxS.

There appear to be subtle and unpredictable influences on occupancy when PNGS are in close proximity, as can be in the case in V1 and, presumably, other variable loops. There are reports that skipping by OST is likely to occur when one or two of the sequons for adjacent PNGS (gap-0 and gap-1) is NxS ^9,19^. In contrast, attachment of glycans by OST is usually efficient when both sequons are NxT. As a result, double NxT sites are highly enriched in libraries of dually glycosylated gap-0 and gap-1 sites derived from mammalian proteins ^9^. That finding is consistent with our observation that glycan attachment at the single gap-0 site in BG505 SOSIP trimers (N190 and N190c) was more efficient when these sequons were changed from NxS-NxS in the WT protein to NxT-NxT in its NxT counterpart.

In contrast, the gap-1 sites N156-N160 and N133-N137 on the BG505 SOSIP trimer did not conform to the above pattern, which was unexpected. In both cases, occupancy was suboptimal when the two sequons were in NxT form, but there was an increase in occupancy when the first site was presented as NxS. One possible explanation is that the x-position of the N156 site is a cysteine that is involved in a disulfide bond, which could interfere with glycan attachment when OST affinity is increased by the NxT change. For both gap-1 sites a lower OST affinity of one of the sequons, whether driven by a lower on-rate and/or a higher off-rate, might allow a different OST complex to more efficiently associate with the neighboring sequon. This kind of competition for OST between sequons might be much more pronounced in glycoproteins with an unusually high glycan density such as HIV-1 Env, compared to more typical human glycoproteins that carry only a few glycans. Hence, we overall conclude that increasing glycan occupancy cannot be achieved by simply changing every sequon to NxT (although doing so often helps); the affinities of sequons for OST and how closely they are spaced are additional factors that need to be taken into account.

The Env trimer is the basis for HIV-1 vaccine research aimed at inducing broad and protective neutralizing antibodies. A now widely used immunogen design platform that is being explored in preclinical and clinical phase tests is the SOSIP trimer, a concept that has evolved from the initial prototype, BG505 SOSIP.664, to improved versions that better resemble viral Env ^47–52^. SOSIP trimers can now be based on most viral sequences from various clades ^14,18,35,37,38,53–57^. However, the SOSIP trimers evaluated to date do not fully mimic viral Env in respect of glycan occupancy; the resulting artificial holes in the glycan shield are immunogenic, but they induce antibodies of generally narrow-specificity that are not on the pathway towards neutralization breadth. Here, we show that the NxT T135S T158S protein closely resembles viral Env in respect of both structure and PNGS occupancy. This next generation Env construct, or others based on the principles derived here, is a suitable starting point for additional immune-focussing efforts aimed at inducing glycan-dependent bNAbs. Finally, the PNGS sequon-engineering strategies we describe here could be applied to other highly glycosylated vaccine antigens and other biologics where PNGS occupancy could usefully be improved.

## Methods

### BG505 infectious molecular clones

The infectious molecular clone of LAI was used as the backbone for creating an infectious molecular clone containing BG505.T332N gp160 ^58,59^. This clone contains a unique Sal1 restriction site 434 nucleotides upstream of the *env* start codon and a unique BamH1 site at the codons specifying amino acids G751 and S752 in LAI gp160 (HXB2 numbering). A DNA fragment containing the LAI sequences between the Sal1 site and the *env* start codon, followed by the BG505.T332N *env* sequences up to the BamH1 site, was synthesized (Genscript, Piscataway, NJ) and cloned into the LAI molecular clone backbone using Sal1 and BamH1. The resulting molecular clone encodes the complete BG505.T332N gp160 sequence, except for the C-terminal 106 amino acids of the cytoplasmic tail, which are derived from LAI gp160. The sequence was verified before the clone was used in virus infectivity and neutralization assays. The resulting virus was able to infect TZM-bl cells and replicate in PBMCs.

The BG505.T332N sequence was modified by mutating all PNGS to either NxT (NxT virus) or NxS (NxS virus). As described above, the resulting *env* sequences, between the Sal1 and BamH1 sites, where obtained from Genscript in a pUC57 cloning vector. The Sal1 and BamH1 BG505-NxT and BG505-NxS sequences were amplified by PCR, using In-Fusion cloning (Clontech), as described by the manufacturer. After the PCR, 0.5 µl DpnI (NEB) was added to each reaction and incubated for 1 h at 37 °C. Next, the PCR products were purified using a PCR clean-up kit (Macherey-Nagel) and the DNA concentrations were measured. The LAI *env* sequences between the Sal1 and BamH1 sites in the pLAI expression plasmid ^59^, were replaced by the BG505-NxT or BG505-NxS PCR fragments using the In-Fusion enzyme (Clontech) as described by the manufacturer. Plasmid DNA (5 µg) of the molecular clones was transfected into HEK293T cells as described previously to generate infectious virus stocks ^38^.

### BG505 SOSIP trimers

The design of BG505 SOSIP.664 and BG505 SOSIP.v4.1 trimers, including the D7324-epitope tagged versions, has been described extensively elsewhere, as have the methods to produce and purify them ^10,35,38,60,61^. BG505 SOSIP.v4.1 expression constructs in which all PNGS were mutated to either NxT (NxT protein) or NxS (NxS protein) were obtained from Genscript and cloned in the pPPI4 expression vector. The Env trimers were purified via PGT145-affinity chromatography and 2G12-affinity chromatography followed by SEC^35,37,38^.

### Antibodies

MAbs were obtained as gifts, or purchased, or expressed from plasmids, from the following sources directly or through the AIDS Reagents Reference Program: John Mascola and Peter Kwong (VRC01); Michel Nussenzweig (3BNC60, 3BC315); Dennis Burton (PG9, PG16, PGT121-123, PGT125-128, PGT130, PGT135, PGT145, PGDM1400 and PGT151); Barton Haynes (CH01); Polymun Scientific (2G12); Mark Connors (35O22); Ms C. Arnold (CA13 (ARP3119)), EU Programme EVA Centralized Facility for AIDS Reagents, NIBSC, UK (AVIP Contract Number LSHP-CT-2004-503487).

### Neutralization assays

One day prior to infection, 1.7 × 10^4^ TZM-bl cells per well were seeded on a 96-well plate in Dulbecco’s Modified Eagles Medium (DMEM) containing 10% FCS, penicillin and streptomycin (both at 100 U/ml), and incubated at 37°C in an atmosphere containing 5% CO_2_. A fixed amount of virus (2.5 ng/ml of p24-antigen equivalent) was incubated for 30 min at room temperature with serial 3-fold dilutions of each test MAb ^38,62,63^. This mixture was added to the cells and 40 µg/ml DEAE, in a total volume of 200 µl. Three days later, the medium was removed. The cells were washed once with PBS (150 mM NaCl, 50 mM sodium phosphate, pH 7.0) and lysed in Lysis Buffer, pH 7.8 (25 mM Glycylglycine (Gly-Gly), 15 mM MgSO_4_, 4 mM EGTA tetrasodium, 10% Triton-X). Luciferase activity was measured using a Bright-Glo kit (Promega, Madison, WI) and a Glomax Luminometer according to the manufacturer’s instructions (Turner BioSystems, Sunnyvale, CA). All infection measurements were performed in quadruple. Uninfected cells were used to correct for background luciferase activity. The infectivity of each mutant without inhibitor was set at 100%. Nonlinear regression curves were determined and 50% inhibitory concentrations (IC_50_) were calculated using a sigmoid function in Prism software version 7.03.

### SDS-PAGE and Blue Native-PAGE

Env proteins were analyzed using SDS-PAGE and BN-PAGE, blotted and detected by using the CA13 (ARP3119) and 2G12 MAbs. In some case the gels were stained using Coomassie blue ^48,64^.

### Negative stain electron microscopy

The WT protein and NxT PNGS mutants were analyzed by negative stain EM as previously described ^21^. For the RM20E1 complexes, SOSIP trimers were incubated with 6-fold molar excess per protomer Fab at RT overnight. The complexes were purified using SEC on a Superose™ 6 Increase 10/300 GL (GE Healthcare) column. Samples were imaged and analyzed as previously described ^21^.

### Cryo electron microscopy

The cryoEM dataset that led to EMDB-21232 was reprocessed with cryoSPARCv.1 to sort particles into classes based on RM20E1 Fab occupancy ^21,65^. Final refinements were performed using cryoSPARCv.1 ^65^.

### D7324-capture ELISA

The D7324-capture ELISA has been described in detail elsewhere ^38^. Microlon 96-well half-area plates (Greiner Bio-One, Alphen aan den Rijn, the Netherlands) were coated with D7324 antibody (10 µg/ml; Aalto Bioreagents, Dublin, Ireland). Purified proteins (2.5 µg/ml) were subsequently captured on the ELISA plate wells and tested for Ab binding. Abs were detected with the goat-anti-human HRP-labeled Ab (Jackson Immunoresearch).

### Biolayer interferometry (BLI)

Antibody binding to the PGT145-purified WT, NxT and NxT T135S T158S PNGS mutants was studied using a ForteBio Octet K2 instrument ^66^. All assays were performed at 30 °C with the agitation setting at 1000 rpm. Purified proteins and antibodies were diluted in running buffer (PBS, 0.1% BSA, 0.02% Tween 20) and analyzed in a final volume of 250 μl/well. Antibody were loaded onto protein A sensors (ForteBio) at 2.0 μg/ml in running buffer until a binding threshold of 0.5 nM was reached. Trimer proteins were diluted in running buffer at 40 nM or 600 nM for CH01, and association and dissociation were measured for 300 s. Trimer binding to a protein A sensor with no loaded antibody was measured to derive background values.

### *N*-glycan analysis using HILIC-UPLC

*N*-linked glycans were released from gp140 in-gel using PNGase F (New England Biolabs). The released glycans were subsequently fluorescently labelled with procainamide using 110mg/ml procainamide and 60mg/ml sodium cyanoborohydride. Excess label and PNGase F were removed using Spe-ed Amide-2 cartridges (Applied Separations). Glycans were analyzed on a Waters Acquity H-Class UPLC instrument with a Glycan BEH Amide column (2.1 mm x 100 mm, 1.7 μM, Waters). Fluorescence was measured, and data were processed using Empower 3 software (Waters, Manchester, UK). The relative abundance of oligomannose glycans was measured by digestion with Endoglycosidase H (Endo H; New England Biolabs). Digestion was performed overnight at 37 degrees. Digested glycans were cleaned using a PVDF protein-binding membrane (Millipore) and analyzed as described above.

### Site-specific glycan analysis using mass spectrometry

Env proteins were denatured for 1h in 50 mM Tris/HCl, pH 8.0 containing 6M of urea and 5mM dithiothreitol (DTT). Next, the Env proteins were reduced and alkylated by adding 20 mM iodacetamide (IAA) and incubated for 1h in the dark, followed by a 1h incubation with 20mM DTT to eliminate residual IAA. The alkylated Env proteins were buffer-exchanged into 50 mM Tris/HCl, pH 8.0 using Vivaspin columns (3 kDa) and digested separately O/N using trypsin, chymotrypsin or subtilisin (Mass Spectrometry Grade, Promega) at a ratio of 1:30 (w/w). The next day, the peptides were dried and extracted using C18 Zip-tip (MerckMilipore). The peptides were dried again, re-suspended in 0.1% formic acid and analyzed by nanoLC-ESI MS with an Easy-nLC 1200 (Thermo Fisher Scientific) system coupled to a Fusion mass spectrometer (Thermo Fisher Scientific) using higher energy collision-induced dissociation (HCD) fragmentation. Peptides were separated using an EasySpray PepMap RSLC C18 column (75 µm × 75 cm). The LC conditions were as follows: 275-minute linear gradient consisting of 0-32% acetonitrile in 0.1% formic acid over 240 minutes followed by 35 minutes of 80% acetonitrile in 0.1% formic acid. The flow rate was set to 200 nL/min. The spray voltage was set to 2.7 kV and the temperature of the heated capillary was set to 40 °C. The ion transfer tube temperature was set to 275 °C. The scan range was 400-1600 m/z. The HCD collision energy was set to 50%, appropriate for fragmentation of glycopeptide ions. Precursor and fragment detection were performed using an Orbitrap at a resolution MS1= 100,000. MS2= 30,000. The AGC target for MS1=4e^5^ and MS2=5e^4^ and injection time: MS1=50ms MS2=54ms

Glycopeptide fragmentation data were extracted from the raw file using Byonic™ (Version 2.7) and Byologic™ software (Version 2.3; Protein Metrics Inc.). The glycopeptide fragmentation data were evaluated manually for each glycopeptide; the peptide was scored as true-positive when the correct b and y fragment ions were observed along with oxonium ions corresponding to the glycan identified. The MS data was searched using a standard library for HEK293F expressed BG505 SOSIP.664 ^67^. The precursor mass tolerance was set at 4ppm for MS1 and 10ppm for MS2. A 1% false discovery rate (FDR) was applied. The relative amounts of each glycan at each site as well as the unoccupied proportion were determined by comparing the extracted ion chromatographic areas for different glycopeptides with an identical peptide sequence. Glycans were categorized according to the composition detected. HexNAc(2)Hex(9–5) was classified as M9 to M5. HexNAc(3)Hex(5–6)X was classified as Hybrid with HexNAc(3)Fuc(1)X classified as Fhybrid. Complex-type glycans were classified according to the number of processed antenna and fucosylation. If all of the following compositions have a fucose they are assigned into the FA categories. HexNAc(3)Hex(3-4)X is assigned as A1, HexNAc(4)X is A2/A1B, HexNAc(5)X is A3/A2B, and HexNAc(6)X is A4/A3B. As this fragmentation method does not provide linkage information isomers are grouped, so for example a triantennary glycan contains HexNAc (5) but so does a biantennary glycan with a bisect.

### Site-specific analysis of low abundance *N*-glycan sites using mass spectrometry

To obtain data for sites that frequently present low intensity glycopeptide the glycans present on the glycopeptides were homogenized to boost the intensity of these peptides. This analysis loses fine processing information but enables the ratio of oligomannose: complex: unoccupied to be determined. The remaining glycopeptides were first digested with Endo H (New England Biolabs) to deplete oligomannose- and hybrid-type glycans and leave a single GlcNAc residue at the corresponding site. The reaction mixture was then dried completely and resuspended in a mixture containing 50 mM ammonium bicarbonate and PNGase F (New England Biolabs) using only H_2_O_18_ (Sigma-Aldrich) throughout. This second reaction cleaves the remaining complex-type glycans but leaves the GlcNAc residues remaining after Endo H cleavage intact. The use of H_2_O_18_ in this reaction enables complex glycan sites to be differentiated from unoccupied glycan sites as the hydrolysis of the glycosidic bond by PNGaseF leaves a heavy oxygen isotope on the resulting aspartic acid residue. The resultant peptides were purified as outlined above and subjected to reverse-phase (RP) nanoLC-MS. Instead of the extensive N-glycan library used above, two modifications were searched for: +203 Da corresponding to a single GlcNAc, a remnant of an oligomannose/hybrid glycan, and +3 Da corresponding to the O^18^ deamidation product of a complex glycan. A lower HCD energy of 27% was used as glycan fragmentation was not required. Data analysis was performed as above and the relative amounts of each glycoform determined, including unoccupied peptides.

## Acknowledgments

We thank J. E. Robinson, J. R. Mascola, P. D. Kwong, D. R. Burton, B. F. Haynes, M. C. Nussenzweig and M. Connors for donating antibodies and reagents directly or through the AIDS Reagents Reference Program. This work was supported by the U.S. National Institutes of Health under grant P0 AI110657 (to J.P.M., A.B.W. and R.W.S.); by the Bill and Melinda Gates Foundation through the Collaboration for AIDS Vaccine Discovery (CAVD) under grants OPP1132237 and INV-002022 (to J.P.M. and R.W.S.), OPP1115782 (to A.B.W.). This work was supported by the IAVI Neutralizing Antibody Center through the CAVD grant OPP1084519 and OPP1196345/INV-008813 funded by the Bill and Melinda Gates Foundation (to M.C. and A.B.W); by the European Union’s Horizon 2020 research and innovation programme under grant agreement No. 681137 (to R.W.S. and M.C.); by the Aids Fonds under grant 2016019 (to R.W.S.); and by the Foundation Dormeur, Vaduz (to R.W.S. and to M.J.v.G.). R.W.S. is a recipient of a Vici grant from the Netherlands Organization for Scientific Research (NWO). M.J.v.G. is a recipient of an AMC Fellowship and a Mathilde Krim Fellowship from the American Foundation for AIDS Research (amfAR) (109514-61-RKVA). C.A.C. is supported by the NIH F31 Ruth L. Kirschstein Predoctoral Award Al131873 and by the Achievement Rewards for College Scientists Foundation. R.D. received a work visit grant from the Amsterdam Infection and Immunity Institute (AI&II). This is manuscript number 30004 from The Scripps Research Institute.

## Contributions

R.D., J.D.A, M.C., R.W.S. designed the study. C.A.C. performed the cryo-EM analysis. J.D.A and R.D performed the overall and site-specific glycan analysis. W-H.L., K.R., A.A., J.C. and S.W.d.T. performed the NS-EM analysis. R.D. and K.S. performed ELISA experiments. R.D., P.W. and T.L.G.M.K performed the virus neutralization assays. A.Y. and P.J.K. performed the surface plasmon resonance analysis. R.D. performed the Biolayer interferometry analysis. M.J.G. isolated the RM20E and RM20E1 MAbs. R.D., J.D.A., P.J.K., G.A., J.P.M., A.B.W., M.C. and R.W.S wrote the manuscript.

## Supplemental Figure Legends

**Table S1.** Occurrence of NxT and NxS sequons in the BG505 sequence compared to 6516 unique Env sequences found in the Los Alamos database. The occupancy of each PNGS on the BG505 SOSIP.v4.1 trimer is also recorded.

|  |  |  | Frequency in natural isolates <sup>1</sup> |  |  |  |
| --- | --- | --- | --- | --- | --- | --- |
| Glycan | BG505 | Motif | % NxT | % NxS | % No PNGS | Glycan occupancy |
| N88 | NxT | NVT | 98.2 | 0.4 | 1.4 | >95% |
| N133 | NxT | NVT | 15.0 | 3.0 | 82.0 | 75% |
| N137 | NxT | NIT | 20.1 | 3.5 | 76.4 | >95% |
| **N156** | **NxS** | **NCS** | **8.6** | **87.6** | **3.8** | **>95%** |
| N160 | NxT | NMT | 88.9 | 2.1 | 9.0 | >95% |
| **N190** | **NxS** | **NRS** | **N.D** <sup>2</sup> | **N.D** <sup>2</sup> | **N.D** <sup>2</sup> | **80%** |
| **N190c** | **NxS** | **NNS** | **N.D** <sup>2</sup> | **N.D** <sup>2</sup> | **N.D** <sup>2</sup> | 75% |
| **N197** | **NxS** | **NTS** | **1.1** | **96.8** | **2.1** | **90%** |
| N234 | NxT | NGT | 74.8 | 5.0 | 20.2 | >95% |
| **N262** | **NxS** | **NGS** | **0.2** | **98.9** | **0.9** | **>95%** |
| N276 | NxT | NIT | 76.5 | 18.5 | 5.1 | >95% |
| N295 | NxT | NCT | 58.2 | 1.5 | 40.4 | >95% |
| N301 | NxT | NNT | 93.7 | 0.2 | 6.2 | >95% |
| **N332** | **NxS** | **NVS** | **3.0** | **68.5** | **28.5** | **>95%** |
| N339 | NxT | NET | 65.1 | 0.1 | 34.8 | >95% |
| N355 | NxT | NNT | 72.8 | 1.6 | 25.5 | >95% |
| **N363** | **NxS** | **NSS** | **0.4** | **7.7** | **91.9** | **>95%** |
| **N386** | **NxS** | **NTS** | **40.0** | **46.8** | **13.2** | **>95%** |
| N392 | NxT | NST | 70.0 | 9.2 | 20.8 | >95% |
| **N398** | **NxS** | **NTS** | **11.0** | **2.9** | **86.1** | **>95%** |
| N406 | NxT | NST | 2.7 | 0.5 | 96.8 | >95% |
| **N411** | **NxS** | **NDS** | **44.4** | **1.3** | **54.3** | **>95%** |
| N448 | NxT | NIT | 87.4 | 0.1 | 12.5 | >95% |
| N463 | NxT | NST | N.D <sup>2</sup> | N.D <sup>2</sup> | N.D <sup>2</sup> | >95% |
| **N611** | **NxS** | **NSS** | **11.6** | **86.8** | **1.6** | **40%** |
| **N618** | **NxS** | **NLS** | **16.6** | **75.3** | **8.1** | **>95%** |
| N625 | NxT | NMT | 96.2 | 0.0 | 3.8 | >95% |
| N637 | NxT | NYT | 92.5 | 3.8 | 3.7 | >95% |

<sup>1</sup> Data obtained from 6516 unique HIV envelope sequences found in the Los Alamos database (http://www.hiv.lanl.gov)
<sup>2</sup> PNGS are located in highly variable regions and the specific sites differ between the natural isolates, making reliable calculations not possible.
Bold indicates the PNGS that have an NxS motif

**Fig. S1.**
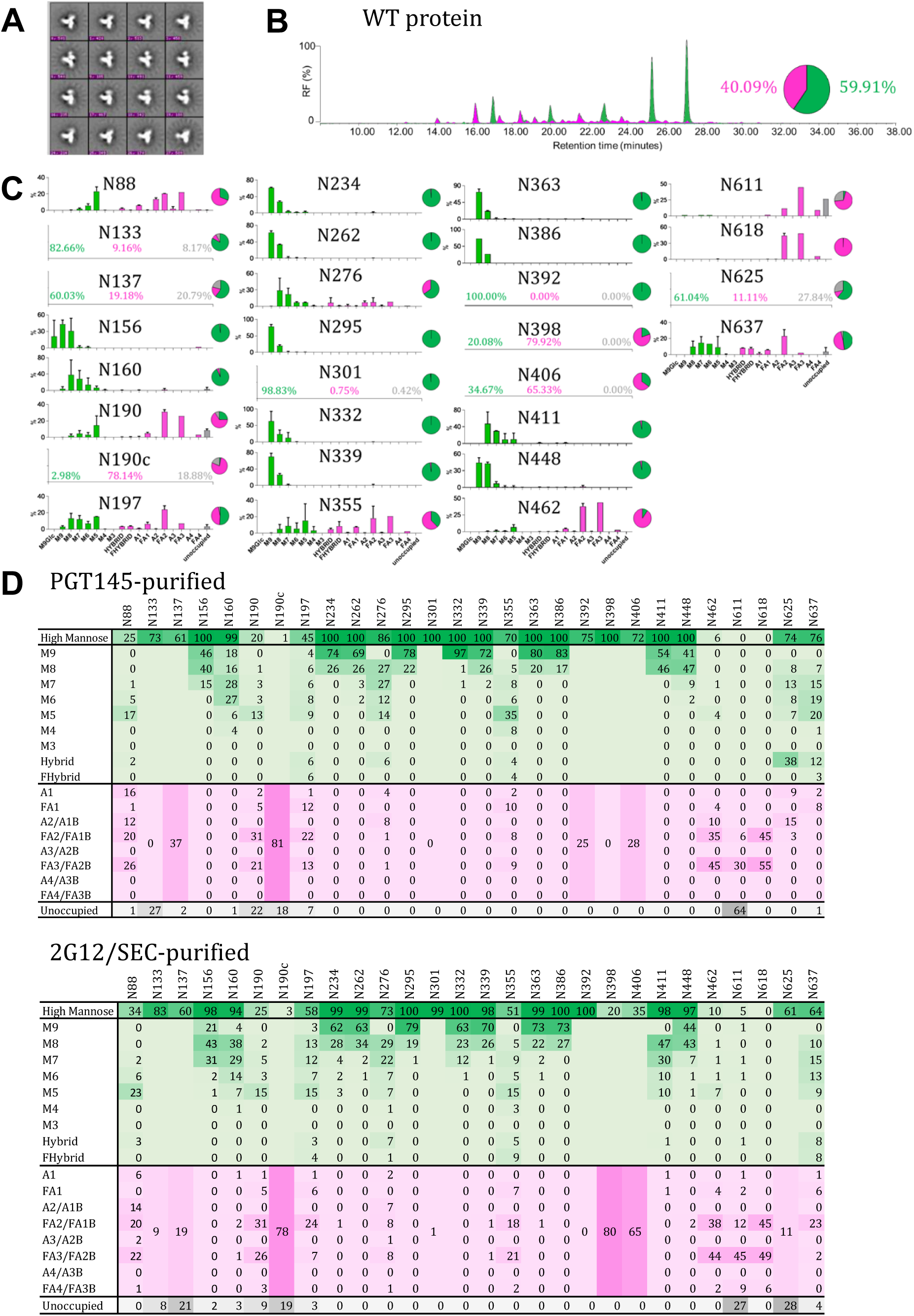
Several PNGS on BG505 SOSIP.v4.1 trimers are underoccupied. **(A)** NS-EM analysis of WT proteins, showing the 2D class-averages. **(B)** HILIC-UPLC analysis of the WT protein. The color coding of the spectra and pie chart is the same as in Fig. 1E **(C)** Quantification of site-specific occupancy and composition for all 28 PNGS on WT proteins, derived from LC-ESI MS experiments. The bar graphs show the glycoforms found at each site. The bar graphs represent the mean of two independent biological replicates of WT protein. The pie charts summarize the relative abundance of oligomannose and complex/hybrid glycans at each individual site, as well as the percentage under-occupancy. **(D)** The data sets show the glycoforms found at each PNGS. Data for oligomannose/hybrid-type glycans are shaded in green, fully processed complex type glycans are shaded in magenta, while the absence of a glycan from some PNGS is shaded in grey. Oligomannose-type glycans are categorized according to the number of mannose residues present, hybrids by the presence/absence of fucose and complex-type glycans by the number of processed antenna and the presence/absence of fucose. For further information see Methods. Data that could only be obtained from low intensity peptides cannot be allocated into the above categories. They are merged to cover all oligomannose/hybrid compositions or complex-type glycans. The data presented in this panel are the mean of two independent biological replicates of WT protein.

**Fig. S2.**
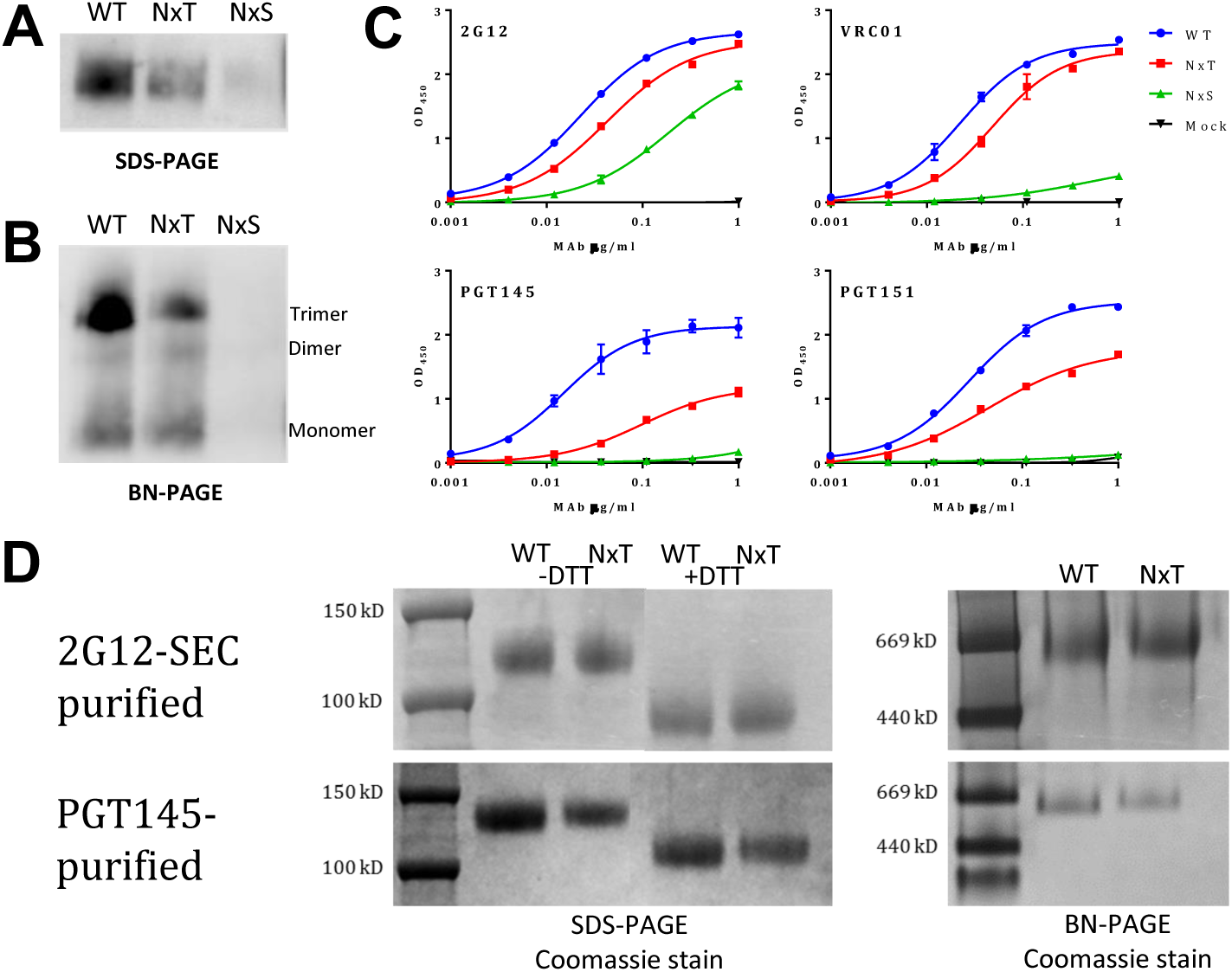
Expression, trimerization and antigenic characterization of WT, NxT and NxS proteins. **(A)** Reducing SDS-PAGE analysis of unpurified WT, NxT and NxS proteins expressed in 293T cells, followed by western blotting with the CA13 (ARP3119) MAb. **(B)** BN-PAGE analysis of the same proteins, blotted with the 2G12 bNAb. The trimer, dimer and monomer bands are indicated. **(C)** D7324-capture ELISA quantifying the binding of the bNAbs 2G12, VRC01, PGT145 and PGT151 to the WT, NxT and NxS proteins. **(D)** Non-reducing and reducing (+ DTT) SDS-PAGE followed by Coomassie staining (left panel), and BN-PAGE analysis followed by Coomassie blue staining (right panel) of PGT145- and 2G12/SEC-purified WT and NxT proteins, as indicated.

**Fig. S3.**
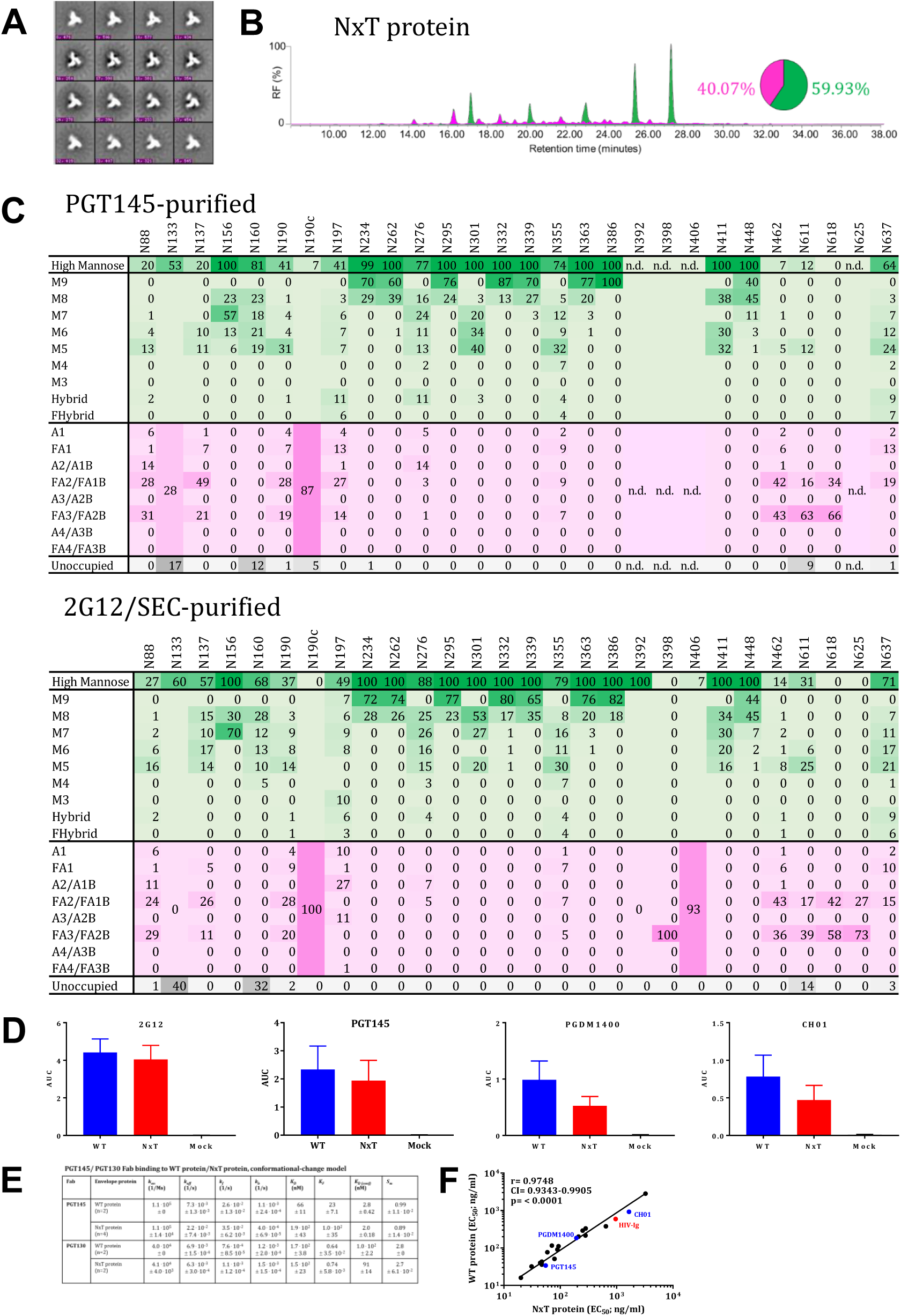
Glycan occupancy is enhanced by PNGS sequon engineering. **(A)** NS-EM analysis of NxT proteins, showing the 2D class-averages. **(B)** HILIC-UPLC analysis of the NxT protein. The color coding of the spectra and pie chart is the same as in Fig. S1B. **(C)** Quantification of site-specific occupancy and composition for 28 PNGS on NxT trimers, purified using the 2G12/SEC and PGT145 methods as indicated. The data are derived from LC-ESI MS experiments. The data set shows the glycoforms found at each PNGS. The relative under-occupancy and oligomannose and complex/hybrid content at each individual site are summarized, using the same color coding as in Fig. S1D. **(D)** Binding of WT and NxT proteins to three V2-apex directed bNAbs, PGT145, PGDM1400 and CH01, and also 2G12 for comparison. AUC values derived from derived from ELISA titration curves are plotted. **(E)** Summary of the SPR binding kinetics of bNAbs PGT145 and PGT130 using data shown in Fig. 2F. **(F)** The EC_50_ values derived using WT and NxT proteins were plotted and compared using Spearman’s correlation coefficient, r, and Prism software version 7.03. Binding data were generated for a panel of bNAbs spanning all major bNAb epitope clusters (Fig. S4). All analyses were performed on NxT proteins produced in HEK293F followed by 2G12/SEC purification.

**Fig. S4.**
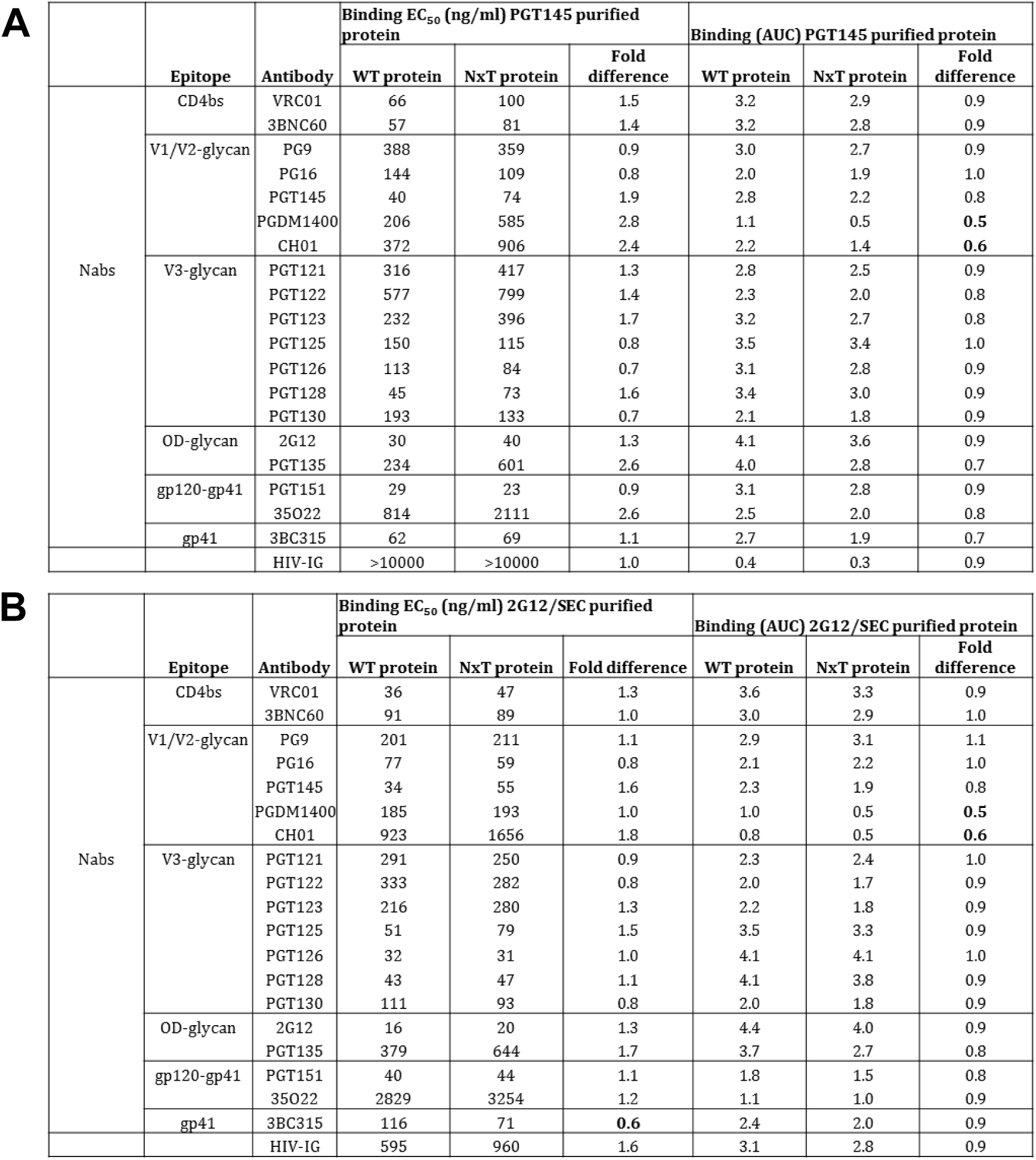
Antigenic characterization of PGT145- and 2G12/SEC-purified WT and NxT proteins. **(A)** Half-maximal binding concentrations (EC_50_; in ng/ml) were derived from D7324-capture ELISAs using PGT145-purified WT and NxT proteins. The values represent the means of 4–10 independent single titration experiments for each bNAb. The fold-differences in EC_50_ values for NxT *vs*. WT proteins are listed. We also tabulated the average AUC values derived from the titration curves for each MAb. Bold numbers indicate values where the differences are <0.6- or >3-fold. **(B)** The same analysis and data presentation but derived using 2G12/SEC-purified WT and NxT proteins.

**Fig. S5.**
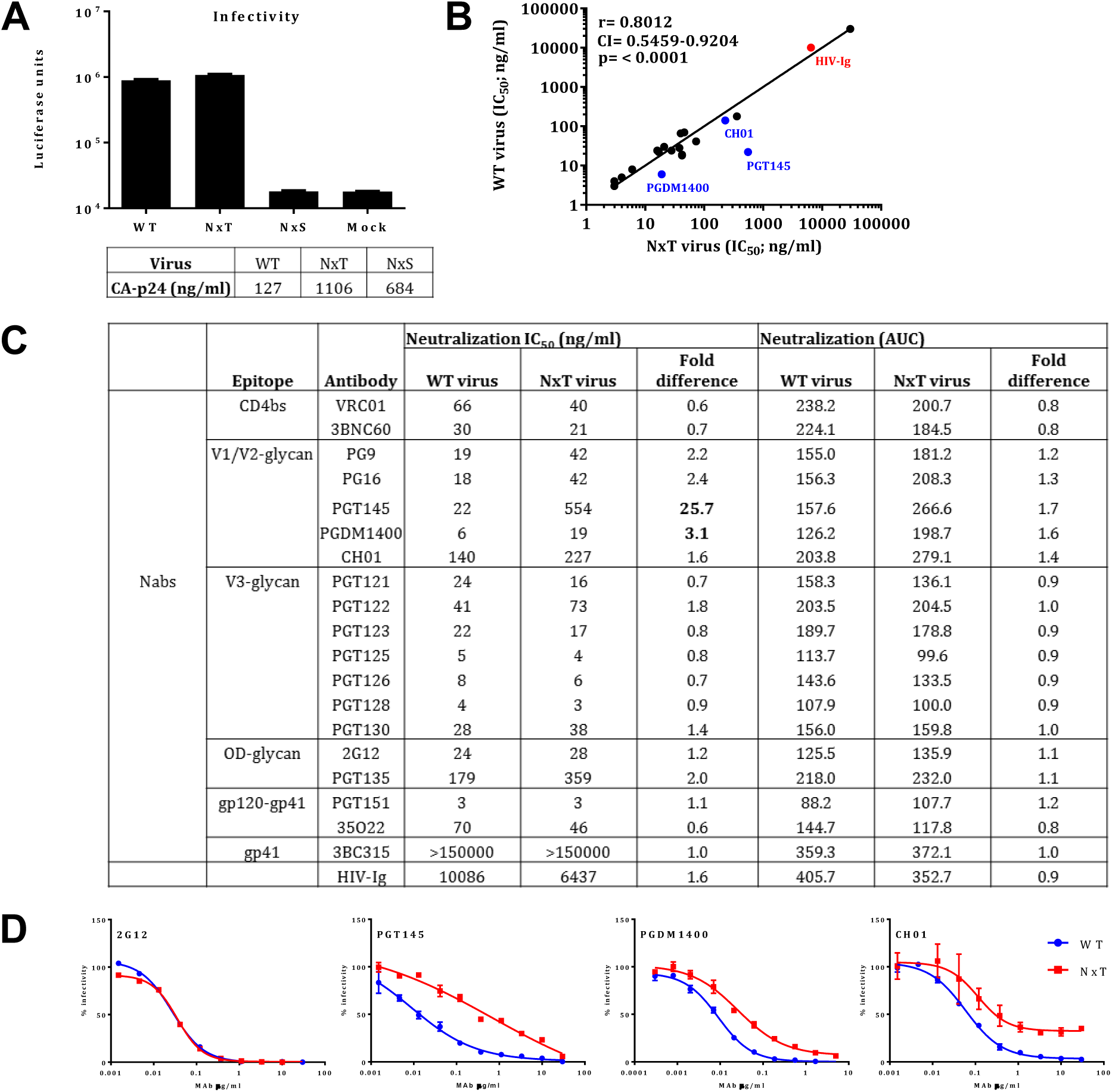
NxT sequon engineering is compatible with Env function and virus infectivity. **(A)** Infection of TZM-bl reporter cells by WT, NxT and NxS viruses. The box represents CA-p24 measured in each virus preparation. **(B)** Correlation between neutralization of WT and NxT viruses. The midpoint neutralization concentrations (IC_50_) were plotted and the Spearman’s correlation coefficient, r, was calculated using Prism software version 7.03. The outliers, PGT145, PGDM1400 and CH01, are indicated in blue. Polyclonal HIV-Ig is indicated in red. **(C)** Midpoint neutralization concentrations (IC_50_; in ng/ml) were derived from single cycle infection experiments using TZM-bl cells and he indicated bNAbs or HIV-Ig. The values are averages based on 2-4 independent antibody-titration experiments. The average AUC values derived from the neutralization curves are also shown, in the three right-most columns. **(D)** Representative neutralization curves for the WT and NxT virus and the 2G12, PGT145, PGDM1400 and CH01 bNAbs.

**Fig. S6.**
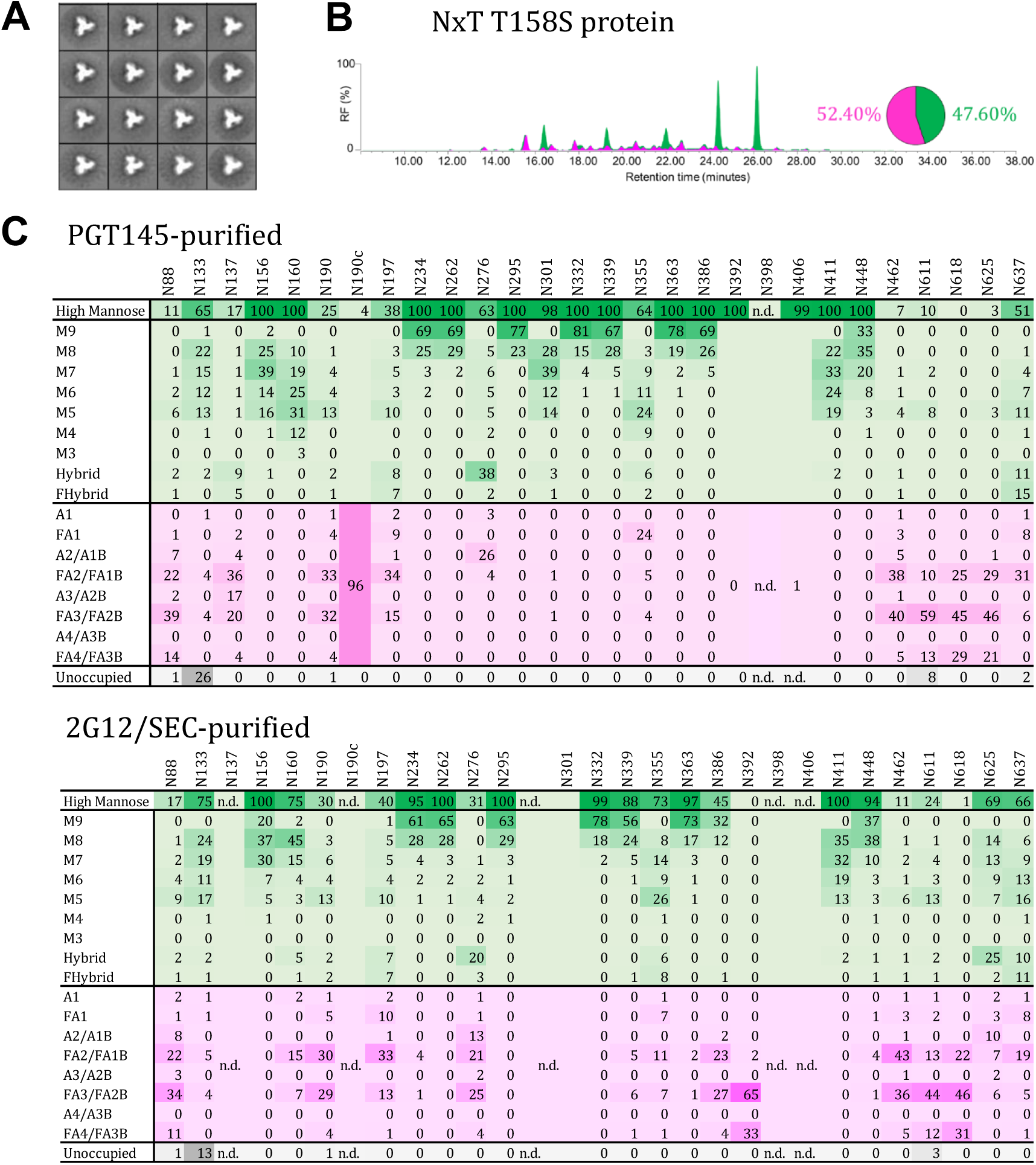
N160 occupancy can be increased by reducing the affinity of a neighboring site for OST. **(A)** NS-EM analysis of NxT 158S trimers, showing the 2D class-averages. **(B)** HILIC-UPLC analysis of the NxT T158S protein. The color coding of the spectra and pie chart is the same as in Fig. S1B. **(C)** Quantification of site-specific occupancy and composition for 28 PNGS on NxT T158S proteins, purified using the 2G12/SEC and PGT145 methods as indicated. The data are derived from LC-ESI MS experiments. The data set shows the glycoforms found at each PNGS. The relative under-occupancy and oligomannose and complex/hybrid content at each individual site are summarized, using the same color coding as in Fig. S1D.

**Fig. S7.**
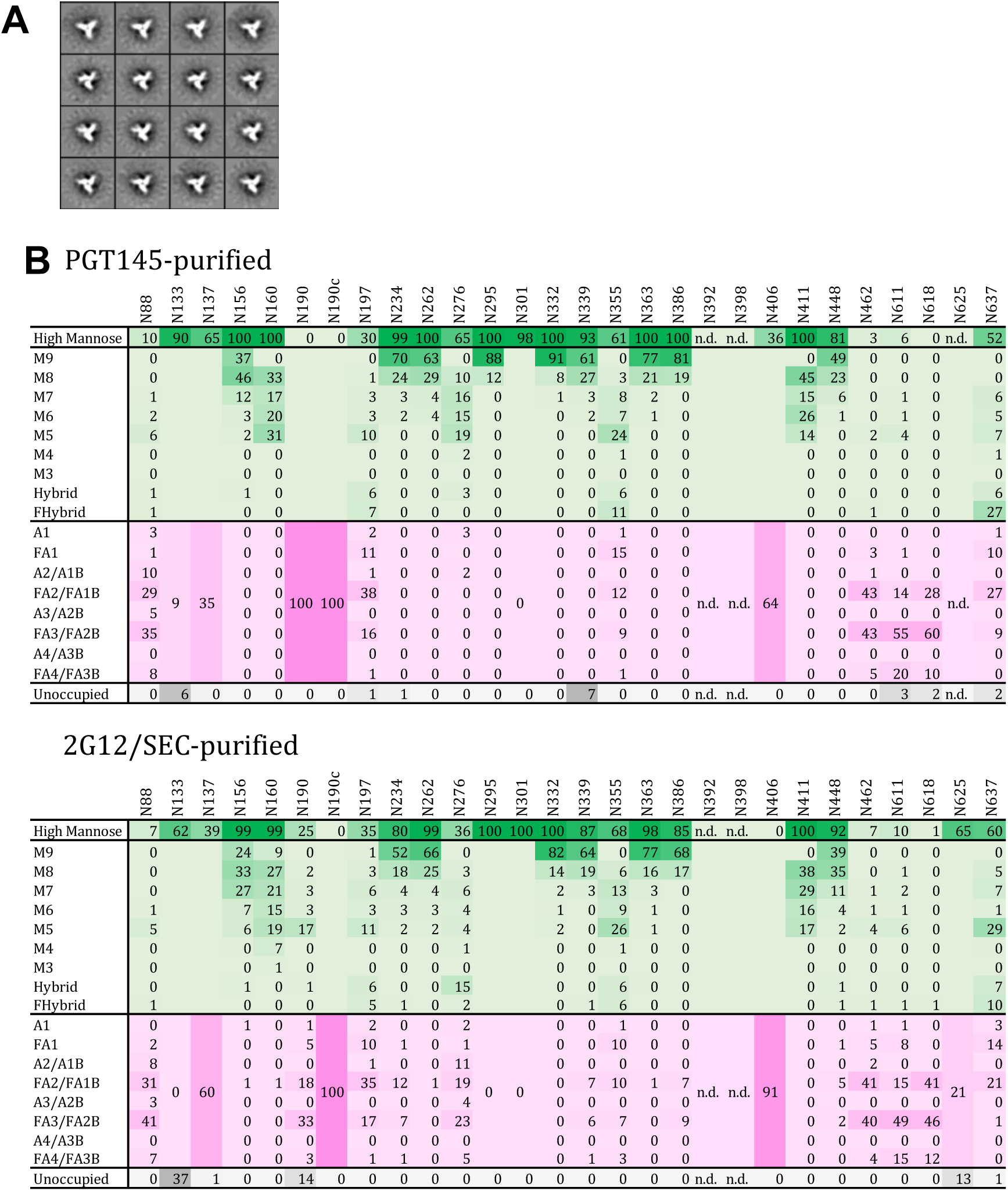
N133 occupancy is increased by reducing its affinity for OST. **(A)** NS-EM analysis of NxT T135S T158S proteins, showing the 2D class-averages. **(B)** Quantification of site-specific occupancy and composition for 28 PNGS on NxT T135S T158S proteins, purified using the 2G12/SEC and PGT145 methods as indicated. The data are derived from LC-ESI MS experiments. The data set shows the glycoforms found at each PNGS. The relative under-occupancy and oligomannose and complex/hybrid content at each individual site are summarized, using the same color coding as in Fig. S1D.

## References

1. Earl, P. L., Moss, B. & Doms, R. W. Folding, interaction with GRP78-BiP, assembly, and transport of the human immunodeficiency virus type 1 envelope protein. J. Virol. 65, 2047–55 (1991).

2. Doms, R. W., Lamb, R. A., Rose, J. K. & Helenius, A. Folding and assembly of viral membrane proteins. Virology 193, 545–62 (1993).

3. Leonard, C. K. et al. Assignment of intrachain disulfide bonds and characterization of potential glycosylation sites of the type 1 recombinant human immunodeficiency virus envelope glycoprotein (gp120) expressed in Chinese hamster ovary cells. J. Biol. Chem. 265, 10373–10382 (1990).

4. Crispin, M., Ward, A. B. & Wilson, I. A. Structure and Immune Recognition of the HIV Glycan Shield. Annu. Rev. Biophys. (2018). doi:10.1146/annurev-biophys-060414-034156

5. Watanabe, Y., Bowden, T. A., Wilson, I. A. & Crispin, M. Exploitation of glycosylation in enveloped virus pathobiology. Biochim. Biophys. Acta – Gen. Subj. 1863, 1480–1497 (2019).

6. Allan, J. S. et al. Major glycoprotein antigens that induce antibodies in AIDS patients are encoded by HTLV-III. Science 228, 1091–1094 (1985).

7. Mellquist, J. L., Kasturi, L., Spitalnik, S. L. & Shakin-Eshleman, S. H. The amino acid following an asn-X-Ser/Thr sequon is an important determinant of N-linked core glycosylation efficiency. Biochemistry 37, 6833–7 (1998).

8. Ruiz-Canada, C., Kelleher, D. J. & Gilmore, R. Cotranslational and posttranslational N- glycosylation of polypeptides by distinct mammalian OST isoforms. Cell 136, 272–83 (2009).

9. Shrimal, S. & Gilmore, R. Glycosylation of closely spaced acceptor sites in human glycoproteins. J. Cell Sci. 126, 5513–23 (2013).

10. Julien, J.-P. et al. Asymmetric recognition of the HIV-1 trimer by broadly neutralizing antibody PG9. Proc. Natl. Acad. Sci. U. S. A. 110, 4351–6 (2013).

11. Sok, D. et al. Recombinant HIV envelope trimer selects for quaternary-dependent antibodies targeting the trimer apex. Proc. Natl. Acad. Sci. U. S. A. 111, 17624–9 (2014).

12. Lee, J. H. et al. A Broadly Neutralizing Antibody Targets the Dynamic HIV Envelope Trimer Apex via a Long, Rigidified, and Anionic β-Hairpin Structure. Immunity 46, 690–702 (2017).

13. Klasse, P. J., Ozorowski, G., Sanders, R. W. & Moore, J. P. Env Exceptionalism: Why Are HIV-1 Env Glycoproteins Atypical Immunogens? Cell Host Microbe 27, 507–518 (2020).

14. Sanders, R. W. et al. HIV-1 neutralizing antibodies induced by native-like envelope trimers. Science 349, (2015).

15. McCoy, L. E. et al. Holes in the Glycan Shield of the Native HIV Envelope Are a Target of Trimer-Elicited Neutralizing Antibodies. Cell Rep. 16, 2327–38 (2016).

16. Wagh, K. et al. Completeness of HIV-1 Envelope Glycan Shield at Transmission Determines Neutralization Breadth. Cell Rep. 25, 893–908.e7 (2018).

17. Ringe, R. P. et al. Closing and opening holes in the glycan shield of HIV-1 envelope glycoprotein SOSIP trimers can redirect the neutralizing antibody response to the newly unmasked epitopes. J. Virol. (2018). doi:10.1128/JVI.01656-18

18. Klasse, P. J. et al. Epitopes for neutralizing antibodies induced by HIV-1 envelope glycoprotein BG505 SOSIP trimers in rabbits and macaques. PLoS Pathog. 14, e1006913 (2018).

19. Cherepanova, N., Shrimal, S. & Gilmore, R. N-linked glycosylation and homeostasis of the endoplasmic reticulum. Curr. Opin. Cell Biol. 41, 57–65 (2016).

20. Struwe, W. B. et al. Site-Specific Glycosylation of Virion-Derived HIV-1 Env Is Mimicked by a Soluble Trimeric Immunogen. Cell Rep. 24, 1958–1966.e5 (2018).

21. Cottrell, C. A. et al. Mapping the immunogenic landscape of near-native HIV-1 envelope trimers in non-human primates. bioRxiv 2020.02.05.936096 (2020). doi:10.1101/2020.02.05.936096

22. Cao, L. et al. Differential processing of HIV envelope glycans on the virus and soluble recombinant trimer. Nat. Commun. 9, 3693 (2018).

23. Kasturi, L., Eshleman, J. R., Wunner, W. H. & Shakin-Eshleman, S. H. The hydroxy amino acid in an Asn-X-Ser/Thr sequon can influence N-linked core glycosylation efficiency and the level of expression of a cell surface glycoprotein. J. Biol. Chem. 270, 14756–61 (1995).

24. Bause, E. Model studies on N-glycosylation of proteins. Biochem. Soc. Trans. 12, 514–7 (1984).

25. Gerber, S. et al. Mechanism of bacterial oligosaccharyltransferase: in vitro quantification of sequon binding and catalysis. J. Biol. Chem. 288, 8849–61 (2013).

26. Pritchard, L. K. et al. Glycan clustering stabilizes the mannose patch of HIV-1 and preserves vulnerability to broadly neutralizing antibodies. Nat. Commun. 6, 7479 (2015).

27. Pritchard, L. K. et al. Structural Constraints Determine the Glycosylation of HIV-1 Envelope Trimers. Cell Rep. 11, 1604–13 (2015).

28. Pritchard, L. K., Harvey, D. J., Bonomelli, C., Crispin, M. & Doores, K. J. Cell- and Protein-Directed Glycosylation of Native Cleaved HIV-1 Envelope. J. Virol. 89, 8932–44 (2015).

29. Bonomelli, C. et al. The glycan shield of HIV is predominantly oligomannose independently of production system or viral clade. PLoS One 6, e23521 (2011).

30. Doores, K. J. et al. Envelope glycans of immunodeficiency virions are almost entirely oligomannose antigens. Proc. Natl. Acad. Sci. U. S. A. 107, 13800–5 (2010).

31. Behrens, A.-J. et al. Composition and Antigenic Effects of Individual Glycan Sites of a Trimeric HIV-1 Envelope Glycoprotein. Cell Rep. 14, 2695–706 (2016).

32. Go, E. P. et al. Comparative Analysis of the Glycosylation Profiles of Membrane- Anchored HIV-1 Envelope Glycoprotein Trimers and Soluble gp140. J. Virol. 89, 8245–57 (2015).

33. Cao, L. et al. Global site-specific N-glycosylation analysis of HIV envelope glycoprotein. Nat. Commun. 8, 14954 (2017).

34. Guttman, M. et al. CD4-induced activation in a soluble HIV-1 Env trimer. Structure 22, 974–84 (2014).

35. de Taeye, S. W. et al. Immunogenicity of Stabilized HIV-1 Envelope Trimers with Reduced Exposure of Non-neutralizing Epitopes. Cell 163, 1702–15 (2015).

36. Torrents de la Peña, A. et al. Improving the Immunogenicity of Native-like HIV-1 Envelope Trimers by Hyperstabilization. Cell Rep. 20, 1805–1817 (2017).

37. Pugach, P. et al. A native-like SOSIP.664 trimer based on a HIV-1 subtype B env gene. J. Virol. 89, 3380–95 (2015).

38. Sanders, R. W. et al. A next-generation cleaved, soluble HIV-1 Env Trimer, BG505 SOSIP.664 gp140, expresses multiple epitopes for broadly neutralizing but not non- neutralizing antibodies. PLoS Pathog. 9, e1003618 (2013).

39. Dey, A. K. et al. cGMP production and analysis of BG505 SOSIP. 664, an extensively glycosylated, trimeric HIV-1 envelope glycoprotein vaccine candidate. Biotechnol. Bioeng. 115, 885–899 (2018).

40. Li, Y., Luo, L., Rasool, N. & Kang, C. Y. Glycosylation is necessary for the correct folding of human immunodeficiency virus gp120 in CD4 binding. J. Virol. 67, 584–588 (1993).

41. Yasmeen, A. et al. Differential binding of neutralizing and non-neutralizing antibodies to native-like soluble HIV-1 Env trimers, uncleaved Env proteins, and monomeric subunits. Retrovirology 11, (2014).

42. Hu, J. K. et al. Murine Antibody Responses to Cleaved Soluble HIV-1 Envelope Trimers Are Highly Restricted in Specificity. J. Virol. 89, 10383–98 (2015).

43. Land, A. & Braakman, I. Folding of the human immunodeficiency virus type 1 envelope glycoprotein in the endoplasmic reticulum. Biochimie 83, 783–90 (2001).

44. Snapp, E. L. et al. Structure and topology around the cleavage site regulate post- translational cleavage of the HIV-1 gp160 signal peptide. Elife 6, (2017).

45. Sellhorn, G., Caldwell, Z., Mineart, C. & Stamatatos, L. Improving the expression of recombinant soluble HIV Envelope glycoproteins using pseudo-stable transient transfection. Vaccine 28, 430–6 (2009).

46. Go, E. P. et al. Glycosylation Benchmark Profile for HIV-1 Envelope Glycoprotein Production Based on Eleven Env Trimers. J. Virol. 91, (2017).

47. Binley, J. M. et al. A recombinant human immunodeficiency virus type 1 envelope glycoprotein complex stabilized by an intermolecular disulfide bond between the gp120 and gp41 subunits is an antigenic mimic of the trimeric virion-associated structure. J. Virol. 74, 627–43 (2000).

48. Sanders, R. W. et al. Stabilization of the soluble, cleaved, trimeric form of the envelope glycoprotein complex of human immunodeficiency virus type 1. J. Virol. 76, 8875–89 (2002).

49. Beddows, S. et al. Construction and characterization of soluble, cleaved, and stabilized trimeric Env proteins based on HIV type 1 Env subtype A. AIDS Res. Hum. Retroviruses 22, 569–79 (2006).

50. Kang, Y. K. et al. Structural and immunogenicity studies of a cleaved, stabilized envelope trimer derived from subtype A HIV-1. Vaccine 27, 5120–32 (2009).

51. Beddows, S. et al. Evaluating the Immunogenicity of a Disulfide-Stabilized, Cleaved, Trimeric Form of the Envelope Glycoprotein Complex of Human Immunodeficiency Virus Type 1 Evaluating the Immunogenicity of a Disulfide-Stabilized, Cleaved, Trimeric Form of the Envelope. (2005). doi:10.1128/JVI.79.14.8812

52. Beddows, S. et al. A comparative immunogenicity study in rabbits of disulfide- stabilized, proteolytically cleaved, soluble trimeric human immunodeficiency virus type 1 gp140, trimeric cleavage-defective gp140 and monomeric gp120. Virology 360, 329–40 (2007).

53. Julien, J.-P. et al. Design and structure of two HIV-1 clade C SOSIP.664 trimers that increase the arsenal of native-like Env immunogens. Proc. Natl. Acad. Sci. U. S. A. 112, 11947–52 (2015).

54. Torrents de la Peña, A. et al. Immunogenicity in Rabbits of HIV-1 SOSIP Trimers from Clades A, B, and C, Given Individually, Sequentially, or in Combination. J. Virol. 92, (2018).

55. Kumar, R. et al. Characterization of a stable HIV-1 B/C recombinant, soluble, and trimeric envelope glycoprotein (Env) highly resistant to CD4-induced conformational changes. J. Biol. Chem. 292, 15849–15858 (2017).

56. Klasse, P. J. et al. Sequential and Simultaneous Immunization of Rabbits with HIV-1 Envelope Glycoprotein SOSIP.664 Trimers from Clades A, B and C. PLoS Pathog. 12, e1005864 (2016).

57. Sliepen, K. et al. Structure and immunogenicity of a stabilized HIV-1 envelope trimer based on a group-M consensus sequence. Nat. Commun. 10, 2355 (2019).

58. Wu, X. et al. Neutralization escape variants of human immunodeficiency virus type 1 are transmitted from mother to infant. J. Virol. 80, 835–44 (2006).

59. Peden, K., Emerman, M. & Montagnier, L. Changes in growth properties on passage in tissue culture of viruses derived from infectious molecular clones of HIV-1LAI, HIV- 1MAL, and HIV-1ELI. Virology 185, 661–72 (1991).

60. Julien, J.-P. et al. Crystal structure of a soluble cleaved HIV-1 envelope trimer. Science 342, 1477–83 (2013).

61. Lyumkis, D. et al. Cryo-EM structure of a fully glycosylated soluble cleaved HIV-1 envelope trimer. Science 342, 1484–90 (2013).

62. Bontjer, I. et al. Optimization of human immunodeficiency virus type 1 envelope glycoproteins with V1/V2 deleted, using virus evolution. J. Virol. 83, 368–83 (2009).

63. Eggink, D. et al. Detailed mechanistic insights into HIV-1 sensitivity to three generations of fusion inhibitors. J. Biol. Chem. 284, 26941–50 (2009).

64. Schülke, N. et al. Oligomeric and conformational properties of a proteolytically mature, disulfide-stabilized human immunodeficiency virus type 1 gp140 envelope glycoprotein. J. Virol. 76, 7760–7776 (2002).

65. Punjani, A., Rubinstein, J. L., Fleet, D. J. & Brubaker, M. A. cryoSPARC: algorithms for rapid unsupervised cryo-EM structure determination. Nat. Methods 14, 290–296 (2017).

66. de Taeye, S. W. et al. Stabilization of the V2 loop improves the presentation of V2 loop-associated broadly neutralizing antibody epitopes on HIV-1 envelope trimers. J. Biol. Chem. 294, 5616–5631 (2019).

67. Seabright, G. E., Doores, K. J., Burton, D. R. & Crispin, M. Protein and Glycan Mimicry in HIV Vaccine Design. J. Mol. Biol. (2019). doi:10.1016/j.jmb.2019.04.016

